# Neuronal glutamate transporters control reciprocal inhibition and gain modulation in D1 medium spiny neurons

**DOI:** 10.1101/2022.07.12.499771

**Authors:** Maurice A. Petroccione, Lianna Y. D’Brant, Nurat Affinnih, Patrick H. Wehrle, Gabrielle C. Todd, Shergil Zahid, Haley E. Chesbro, Ian L. Tschang, Annalisa Scimemi

**Author notes:** Corresponding author and lead contact: Annalisa Scimemi, PhD, SUNY Albany, Department of Biology, 1400 Washington Avenue, Albany (NY) 12222-0100, USA, or.

## Abstract

Understanding the function of glutamate transporters has broad implications for explaining how neurons integrate information and relay it through complex neuronal circuits. Most of what is currently known about glutamate transporters, specifically their ability to maintain glutamate homeostasis and limit glutamate diffusion away from the synaptic cleft, is based on studies of *glial* glutamate transporters. By contrast, little is known about the functional implications of *neuronal* glutamate transporters. The neuronal glutamate transporter EAAC1 is widely expressed throughout the brain, particularly in the striatum, the primary input nucleus of the basal ganglia, a region implicated with movement execution and reward. Here, we show that EAAC1 limits synaptic excitation onto a population of striatal medium spiny neurons identified for their expression of D1 dopamine receptors (D1-MSNs). In these cells, EAAC1 also contributes to strengthen lateral inhibition from other D1-MSNs. Together, these effects contribute to reduce the gain of the input-output relationship and increase the offset at increasing levels of synaptic inhibition in D1-MSNs. By reducing the sensitivity and dynamic range of action potential firing in D1-MSNs, EAAC1 limits the propensity of mice to rapidly switch between behaviors associated with different reward probabilities.

## Introduction

The *neuronal* glutamate transporter EAAC1 is distributed broadly throughout the brain (Rothstein et al. 1994; Shashidharan et al. 1997), with plasma membrane surface density values thought to be significantly lower than those of *astrocytic* glutamate transporters (Holmseth et al. 2012). Glutamate binding to glutamate transporters activates currents that can be easily recorded in heterologous expression systems and astrocytes, where the density of expression of these molecules is high (Wadiche, Amara, and Kavanaugh 1995). In most neurons, recording glutamate transporter mediated currents continues to be elusive (Holmseth et al. 2012). This finding sparked some doubts on the functional relevance of neuronal glutamate transporters like EAAC1 in the brain (Holmseth et al. 2012). Despite this concern, multiple lines of evidence point to the fact that EAAC1 is an important player in the regulation of synaptic function and behavior. For example, in the hippocampus, EAAC1 limits NMDA receptor activation and increases GABA release (Diamond 2001; Scimemi, Tian, and Diamond 2009; Mathews and Diamond 2003). In the dorsolateral striatum (DLS), EAAC1 limits activation of group I metabotropic glutamate receptors (mGluRI) and is associated with increased execution of stereotyped motor behaviors (Bellini et al. 2018). While most of these works relied on the use of EAAC1^-/-^ mice (Peghini, Janzen, and Stoffel 1997), other studies that used overexpression models of EAAC1 showed behavioral abnormalities, including increased anxiety-like behaviors (Delgado-Acevedo et al. 2019; Escobar et al. 2021) and reduced responses to amphetamine-induced hyperlocomotion (Zike et al. 2017). The emerging picture is that maintaining optimal levels of EAAC1 expression is key for proper execution of a wide range of complex motor behaviors and habitual actions, many of which are critically dependent on the activity of the DLS (Gremel and Costa 2013; Yin, Knowlton, and Balleine 2006, 2004).

What continues to puzzle the field is how we can reconcile the obvious behavioral abnormalities of EAAC1^-/-^ mice with the apparent inability to record EAAC1-mediated currents in neurons. A possible way out of this conundrum comes from considerations of the biophysical properties and sub-cellular distribution of this neuronal transporter. *First,* EAAC1 has a very low single channel conductance (∼0.3 fS) (Grewer et al. 2000) and is not evenly distributed along the plasma membrane (Cheng et al. 2002; Conti et al. 1998; Rothstein et al. 1994; He et al. 2000). *Second,* the fact that currents generated at a distance from the soma are dramatically reduced by electrotonic filtering and attenuation, could make them particularly challenging to identify using somatic patch-clamp recordings (Tonnesen et al. 2014; Svoboda, Tank, and Denk 1996; Rall 1959). *Third*, tentative estimates of the *average* surface density of expression EAAC1 obtained from immunoblot experiments do not provide information on the *local* density of expression of EAAC1 in dendritic spines and axonal boutons, where EAAC1 is thought to be confined. *Fourth*, the amplitude of local EAAC1-mediated currents does not provide a direct readout of the number of receptors that EAAC1 might protect from glutamate spillover (Scimemi, Tian, and Diamond 2009). Together, these findings suggest that the inability to record EAAC1-mediated currents from the soma should not be interpreted as evidence of a lack of any functional role for glutamate uptake via EAAC1.

Interestingly, qualitative pre-, and post-embedding ultrastructural works show that EAAC1 has a distinctive punctate peri-synaptic and post-synaptic expression in glutamatergic neurons, attributed to the presence of a specific domain in the C-terminal region that controls the targeting of EAAC1 to dendritic spines and shafts (Cheng et al. 2002; He et al. 2000). EAAC1 is also expressed in a population of GABAergic neurons identified by their expression of glutamic acid decarboxylase, the biosynthetic enzyme for GABA (Conti et al. 1998). In these neurons, EAAC1 shows a clustered expression in a subset of axonal presynaptic terminals (He et al. 2000; Rothstein et al. 1994), although these findings have been brought into question (Holmseth et al. 2012). In this case, the inconsistent results may be due to the limited sensitivity and specificity of antibodies directed against EAAC1 used for immunolabeling studies, which is a historically challenging, if not insurmountable issue. Other labeling strategies based on the use of pharmacological tools have been hampered by the fact that there continues to be no drug that targets EAAC1 without also affecting other glial glutamate transporters (Shimamoto et al. 1998; Tsukada et al. 2005). For these reasons, functional studies combined with genetic approaches currently represent an ideal tool to fill current gaps of knowledge on the functional roles of EAAC1 on synaptic communication in different brain areas.

Multiple lines of evidence suggest the existence of potential association between loss of function of EAAC1, altered synaptic activity in the DLS, and obsessive-compulsive disorder (OCD). *First,* the striatum is one of the brain regions with the most abundant expression of EAAC1. *Second,* hyperactivity of striatal circuits is associated with the emergence of compulsive behaviors in OCD (Gasso et al. 2015; Gilbert et al. 2008; Menzies, Williams, et al. 2008; Abramowitz, Taylor, and McKay 2009; Carmin et al. 2002). *Third,* several genome-wide studies identified loss-of-function variants and single nucleotide polymorphisms of *SLC1A1*, the homolog gene encoding EAAC1 in humans diagnosed with OCD (Veenstra-VanderWeele et al. 2012; Wendland et al. 2009; Porton et al. 2013; Stewart et al. 2007; Stewart, Mayerfeld, et al. 2013; Stewart, Yu, et al. 2013). *Fourth,* one of the key hallmarks of OCD is disrupted balance between goal-directed and habitual actions, both of which fall under the control of striatal circuits (Gillan et al. 2011).

One of the hypotheses that has been brought forward to explain how EAAC1 may limit striatal hyperactivity is that impairing glutamate uptake via EAAC1 might lead to an increased ambient glutamate concentration in the striatum, which in turn would trigger hyperactivity (Porton et al. 2013). This interpretation is difficult to reconcile with evidence that EAAC1 contributes only ∼5% of all glutamate transporters (Holmseth et al. 2012). Even if all EAAC1 molecules lost their ability to bind and transport glutamate, under steady-state conditions, the remaining ∼95% of glutamate transporters in glial cells would be able to keep the extracellular glutamate concentration at low nanomolar levels (Herman and Jahr 2007; Chiu and Jahr 2017). Accordingly, experimental data show that loss of EAAC1 does not change the extracellular glutamate concentration (Rothstein et al. 1996), a result supported also by our own previous works in the striatum (Bellini et al. 2018) and hippocampus (Scimemi, Tian, and Diamond 2009). An alternative hypothesis comes from computational models of neural dynamics in cortico-striatal-thalamic networks, which suggest that loss of EAAC1 might contribute to striatal hyperactivity by changing the relative strength of synaptic excitation and inhibition (E/I) locally, in some or all medium spiny neurons (MSNs), the main long-range projection neurons in the striatum (Rǎdulescu et al. 2017). According to this model, local changes in E/I of striatal MSNs can alter the firing rates of neurons not only in the striatum but also in larger neural networks that include the cortex and thalamus, bringing them in a regime of hyperactivity (Rǎdulescu et al. 2017). Consistent with these findings, abnormal firing rates in MSNs occur in the caudate nucleus of OCD patients during the manifestation of their symptoms (Guehl et al. 2008; Burguiere et al. 2015). Although these theoretical inferences provide potential explanations of how local changes in E/I may lead to changes in the activity of more complex neural networks and behaviors, an experimental investigation of how EAAC1 alters E/I in different populations of striatal MSNs has not been previously performed.

Here, we show that EAAC1 limits synaptic excitation and strengthens synaptic inhibition in the DLS (i.e., reduces E/I), by modulating the strength of excitatory synaptic transmission onto D1 dopamine receptor expressing MSNs (D1-MSNs), and reciprocal inhibition among D1-MSNs. Through these mechanisms, EAAC1 increases the offset of the input-output relationship of D1-MSNs, to levels that differ depending on the rate of incoming inhibition. Together, these findings indicate that impairing the activity of EAAC1, as it might happen in OCD, brings D1-MSNs in a hyperexcitable state where these cells fire more action potentials in response to *(i)* lower frequencies and *(ii)* smaller changes in the frequency of activation of thalamo-cortical excitatory inputs. The reduced lateral inhibition among D1-MSNs may promote task switching, increasing the likelihood that mice rapidly change from one motor behavior to another. These findings shed new light on the synaptic and circuit mechanisms through which EAAC1 modulates the activity of specific populations of MSNs, with potential implications for understanding the cellular basis of impulsivity and the control of executive function in the broad context of OCD (Grassi et al. 2015; Bari and Robbins 2013; Boisseau et al. 2012; Benatti et al. 2014; Ettelt et al. 2007).

## Results

### EAAC1 is expressed in DLS MSNs

Although there is evidence that EAAC1 is expressed in the DLS (Holmseth et al. 2012), it is not known whether its cellular distribution is limited to MSNs or differs between D1- and D2-MSNs. To address this, we performed an RNAscope FISH analysis in the DLS using probes for *Drd1a*, *Drd2* and *Slc1a1*, the genes encoding D1/2 dopamine receptors and EAAC1 in mice, respectively (**Fig. 1A, B**). We identified a population of cells that did not express EAAC1, which we excluded from further analysis (24% of all cells stained with the nuclear marker DAPI; **Fig. 1B**). The EAAC1-expressing cells could be further classified into three main groups using unsupervised clustering (**Fig. 1C**), principal component analysis (**Fig. 1D**) and a dimensionality reduction approach based on t-distributed stochastic neighbor embedding (t-SNE) analysis (**Fig. 1F**). These three EAAC1-expressing cohorts of cells were identified as being either D1-MSNs (25%), D2-MSNs (24%; **Fig. 1E**), or cells with a very low, but detectable expression of EAAC1 and D1/2 receptors (51%; **Fig. 1E**). These findings suggest that: *(i)* in the DLS, EAAC1 mRNA is mostly expressed in MSNs compared to other types of cells, and *(ii)* at the mRNA level, the expression of EAAC1 is indistinguishable between D1- and D2-MSNs (**Fig. 1G, H**).

**Figure 1.**
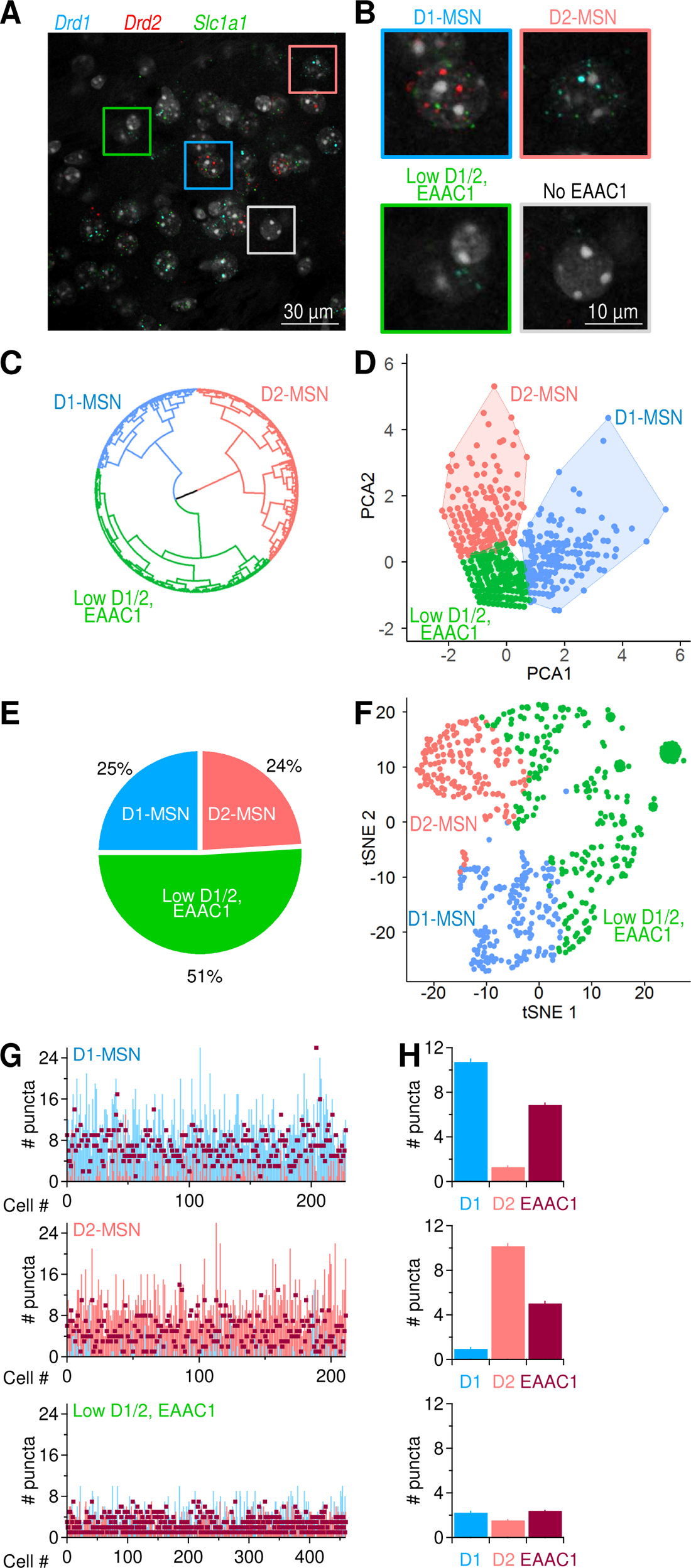
Distribution of EAAC1 mRNA in the DLS. **(A)** RNAscope FISH images of *Drd1* (*blue*), *Drd2* (*red*) and *Slc1a1* (*green*) mRNA expression in the DLS. These transcripts encode D1/2 receptors and EAAC1, respectively. **(B)** Annotated cell clusters identified with color-coded boxes in A, including cells expressing *Drd1* (i.e., D1-MSNs; *blue*), cells expressing *Drd2* (i.e., D2-MSNs; *salmon*), cells with low *Drd1/Drd2/ Slc1a1* EAAC1 expression (*green*), and cells with no detectable *Slc1a1* expression (which also had a very low expression of *Drd1/Drd2*; *gray*). **(C)** Circular dendrogram representation of RNAscope FISH data from EAAC1-expressing cells in the DLS. The dendrogram was obtained using the Ward’s minimum variance method for hierarchical cluster analysis, using three as the number of groups cutting the tree (i.e., the optimal number of clusters obtained using gap statistics). The three main groups represent D1-MSNs (*blue*), D2-MSNs (*salmon*) and cells that express low levels of D1, D2 and EAAC1 (*green*). **(D)** Principal component analysis obtained using an enhanced k-means clustering with the three populations of cells. **(E)** Pie chart describing the percentage of the cell types identified with the hierarchical clustering. **(F)** Annotated clusters in the t-SNE map showing three specific groups of cells identified by the k-means algorithm based on their differential expression of D1, D2 and EAAC1. **(G)** Analysis of mRNA puncta in D1-MSNs (n=229; *blue*), D2-MSNs (n=212, *salmon*) and low D1/2, EAAC1 (n=460, *burgundy*). **(H)** Summary graph showing the mean number of puncta measured in each of the three identified cell populations described in C.

### EAAC1 limits dendritic branching in D1-MSNs

There is ample evidence that the morphology of neurons is susceptible to changes in excitatory synaptic function and connectivity. This has been observed across multiple brain regions, including the visual system, where the length and number of dendrites in stellate cells is reduced by visual deprivation, and the cerebellum, where Purkinje neurons have stunted dendrites when they receive fewer synapses from granule cells (Coleman and Riesen 1968; Wiesel and Hubel 1963; Guillery 1973; Friedlander, Stanford, and Sherman 1982; McAllister 2000; Rakic 1975; Rakic and Sidman 1973; Sotelo 1975). Based on this, we reasoned that a structural analysis of MSNs could provide insights into potential functional changes in synaptic transmission associated with loss of EAAC1 expression. We reconstructed tdTomato-expressing and biocytin-filled MSNs from the DLS of mice that either expressed EAAC1 (i.e., D1^Cre/+^:Ai9^Tg/Tg^:EAAC1^+/+^ and A2A^Cre/+^:Ai9^Tg/Tg^:EAAC1^+/+^ mice) or constitutively lacked it (i.e., D1^Cre/+^:Ai9^Tg/Tg^:EAAC1^-/-^ and A2A^Cre/+^:Ai9^Tg/Tg^:EAAC1^-/-^ mice; **Fig. 2A, B**). For simplicity, in the remainder of this manuscript, we collectively refer to all these mice as EAAC1^+/+^ and EAAC1^-/-^, respectively. A morphological Sholl analysis showed that D1-MSNs span a larger territory of the DLS neuropil and have a more intricate dendritic architecture in EAAC1^-/-^ (n=10) compared to EAAC1^+/+^ mice (n=13; ***p=9.4e-6, r=0.90; **Fig. 2C, *left***). This effect that was not detected when comparing D2-MSNs in the two mouse strains (EAAC1^+/+^ mice, n=10; EAAC1^-/-^, n=11, p=0.70, r=0.94; **Fig. 2C, *right***). Consistent with these findings, several other measures confirmed that the dendritic arbor of D1-but not D2-MSNs was larger in the absence of EAAC1. *First,* the maximum radius of the dendritic field of D1-MSNs was larger in EAAC1^-/-^ mice (*p=0.02), whereas that of D2-MSNs was similar in EAAC1^+/+^ and EAAC1^-/-^ mice (p=0.09; **Fig. 2D, *left***). *Second,* the center of mass of the dendritic arbor was located further away from the soma in D1-MSNs of EAAC1^-/-^ compared to EAAC1^+/+^ mice, whereas this was not the case for D2-MSNs (D1-MSN: **p=7.6e-3; D2-MSN: p=0.17; **Fig. 2D, *center***). *Third,* the area of the neuropil covered by D1-MSNs was 2-fold larger in D1-MSNs of EAAC1^-/-^ compared to EAAC1^+/+^ mice (*p=0.02), but it was similar in D2-MSNs (p=0.54; **Fig. 2D, *right***). *Fourth,* the total length of all dendrites, which receive synaptic inputs from other neurons, was ∼2-fold larger in D1-MSNs of EAAC1^-/-^ compared to EAAC1^+/+^ mice (*p=0.011), but similar in D2-MSNs (p=0.27; **Fig. 2E**). *Fifth,* the dendritic arbor of D1-MSNs in EAAC1^-/-^ versus EAAC1^+/+^ mice had more branches (*p=0.03; **Fig. 2F, *left***) and branching points (*p=0.03; **Fig. 2F, *center***), with a similar length of branch segments between branching points (p=0.70; **Fig. 2F, *right***). Dendrites of D2-MSNs had the same number of branches (p=0.18; **Fig. 2F, *left***), branching points (p=0.19; **Fig. 2F, *center***), and branch segment length (p=0.51; **Fig. 2F, *right***). These findings indicate that, through mechanisms that remain to be determined, EAAC1 is implicated with regulating the dendritic architecture of D1-MSNs.

**Figure 2.**
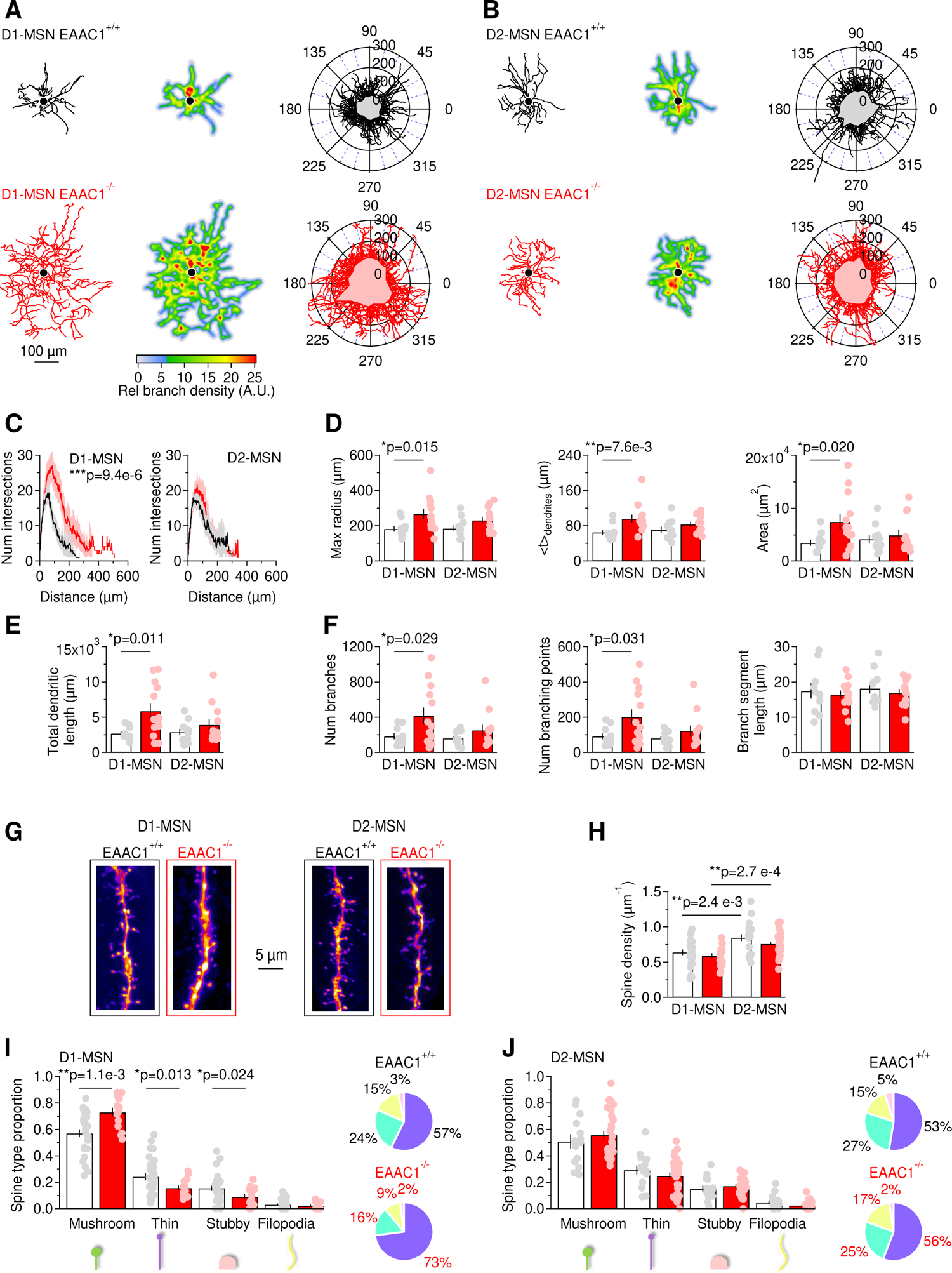
EAAC1 limits the branching pattern and spine size in D1-MSNs. **(A)** *Left,* Example of biocytin-filled and reconstructed D1-MSNs in EAAC1^+/+^ (*top, black*) and EAAC1^-/-^ mice (*bottom, red*). *Center,* heat maps of the reconstructions shown in the left panel. Regions highlighted in red are the areas with the highest branch density. *Right,* polar plots showing overlaid reconstructions of D1-MSNs in EAAC1^+/+^ (n=10) and EAAC1^-/-^ mice (n=13). The gray and pink shaded areas represent the mean coverage area of D1-MSNs in each mouse genotype. **(B)** As in **A**, for D2-MSNs in EAAC1^+/+^ (n=11) and EAAC1^-/-^ mice (n=12). **(C)** *Left,* Sholl plots showing the number of dendritic branch intersections formed by D1-MSNs at increasing distances from the center of the soma, for the D1-MSNs described in A. *Right,* As in left, for D2-MSNs. **(D)** *Left,* The maximum radius of dendritic branches, calculated as the maximum x-value in the Sholl plots in C, is increased in D1-MSNs of EAAC1^-/-^ mice. *Center,* The geometric centroid (*<t>*) is the center of mass of the Sholl plots, which is the distance from the soma before and after which there is the same density of branches (i.e., number/weight). *Right,* Analysis of DLS neuropil coverage by MSNs, showing that D1-MSNs occupy a larger domain of the DLS neuropil in EAAC1^-/-^ mice. **(E)** The total length of all dendrites is larger in D1-MSNs of EAAC1^-/-^ mice. **(F)** Dendritic branch analysis of DLS MSNs. The total number of dendritic branches (*left*) and the number of branching points (*center*) are increased in D1-MSNs of EAAC1^-/-^ compared to EAAC1^+/+^ mice. The mean branch length is similar among MSNs of EAAC1^+/+^ and EAAC1^-/-^ mice. **(G)** 2D maximum intensity projections of confocal images of dendritic branches from D1-MSNs (*left*) and D2-MSNs (*right*) in EAAC1^+/+^ (*black frame*) and EAAC1^-/-^ mice (*red frame*). **(H)** The density of dendritic spines is larger in D2-MSNs compared to D1-MSNs, in EAAC1^+/+^ and EAAC1^-/-^ mice (EAAC1^+/+^ D1-MSNs n=27, D2-MSNs n=17; EAAC1^-/-^ D1-MSN n=18, D2-MSN n=38). **(I)** Spine classification analysis, showing that there is an increase in the proportion of mushroom spines and a decrease in the proportion of thin and stubby spines in D1-MSNs from EAAC1^-/-^ mice. The pie charts on the right show the distribution of different spine types in D1-MSNs from EAAC1^+/+^ (*black*) and EAAC1^-/-^ mice (*red*). Same n values as in **H**. **(J)** As in I, for D2-MSNs. Different types of spines are equally represented in D2-MSNs from EAAC1^+/+^ and EAAC1^-/-^ mice. Same n values as in H. Data represent mean ± SEM.

### EAAC1 reduces spine number and size in D1-MSNs

Dendritic growth and an increased complexity of the dendritic arbor can be promoted by increased neuronal activity, *in vitro* and *in vivo* (Yu and Malenka 2003; Redmond, Kashani, and Ghosh 2002; Sin et al. 2002). Since changes in dendritic morphology are often concurrent with changes in synaptic structure and function, we asked whether the structural changes in the dendritic arborization of D1-MSNs in the absence of EAAC1 might be associated with changes in the number and functional properties of excitatory synaptic inputs onto these cells (Peng et al. 2009). We addressed this question by analyzing the density and subtype distribution of dendritic spines, the anatomical correlates of excitatory synapses, in biocytin-filled MSNs. Spines size scales with their AMPA receptor content and the release probability of the presynaptic terminal with which they are in contact (Schikorski and Stevens 1997). We found that the spine density along the dendrites of D1-MSNs and D2-MSNs was not altered in the absence of EAAC1 (p=0.34 and p=0.12, respectively), but was higher in D2-compared to D1-MSNs in EAAC1^+/+^ and EAAC1^-/-^ mice (**p=2.4e-3 and **p=2.7e-4, respectively; **Fig. 2G, H**). Given that the total dendritic length is larger in EAAC1^-/-^ mice, the presence of a similar density of spines in D1-MSNs of EAAC1^+/+^ and EAAC1^-/-^ mice indicates that D1-MSNs receive a larger number of excitatory synaptic inputs in the absence of EAAC1 (spine number = spine density · dendritic length). In addition, there was an increase in the proportion of mushroom spines and a decrease in that of thin and stubby spines in D1-MSNs of EAAC1^-/-^ mice (D1-MSN mushroom spines: **p=1.1e-3; thin spines: *p=0.01; stubby spines: *p=0.02; **Fig. 2I**). This effect was not detected when comparing the spine size distribution in D2-MSNs of EAAC1^+/+^ and EAAC1^-/-^ mice (D2-MSN mushroom spines: p=0.28; thin spines: p=0.15; stubby spines: p=0.37; **Fig. 2J**). Therefore, by reducing the size of the dendritic arbor of D1-MSNs, EAAC1 allows these cells to receive fewer and weaker excitatory synaptic inputs.

### EAAC1 limits excitation onto D1-MSNs

An increase in the spine number and size in D1-MSNs of EAAC1^-/-^ mice might be indicative of functional changes in the quantal parameters (*N, p* and *q*), which can manifest themselves as changes in the frequency and/or kinetics of action potential-independent miniature EPSCs (mEPSCs). Therefore, we recorded mEPSCs in the presence of the voltage-gated sodium channel blocker tetrodotoxin (TTX 1 µM) and analyzed their frequency, amplitude and kinetics. In our experiments, the mEPSC frequency was larger in D1-MSNs of EAAC1^-/-^ mice compared to EAAC1^+/+^ mice (*p=0.01; **Fig. 3A, B**). This result is consistent with an increase in the number and/or release probability of excitatory synaptic contacts onto D1-MSNs in EAAC1^-/-^ mice. To determine whether the increased mEPSC frequency in D1-MSNs of EAAC1^-/-^ mice could be attributed exclusively to differences in *N* (the number of functional release sites) or also differences in *p* (the release probability), we measured the paired-pulse ratio (PPR) of evoked AMPA EPSCs in D1-MSNs, as this parameter is inversely proportional to the release probability, *p*. The PPR of AMPA EPSCs was similar in D1-MSNs from EAAC1^-/-^ and EAAC1^+/+^ mice (p=0.43), arguing against differences in *p* between excitatory synapses onto D1-MSNs in EAAC1^-/-^ and EAAC1^+/+^ mice. The mEPSC analysis also revealed that the mEPSC amplitude was larger in the absence of EAAC1 (**p=1.8e-3; **Fig. 3C**), with no significant difference in mEPSC kinetics (rise: p=0.87; t_50_: p=0.82; **Fig. 3D**). This is consistent with an increased quantal size (*q*) at excitatory synapses onto D1-MSNs in EAAC1^-/-^ mice, and with the increased spine size in these cells. Therefore, both *N* and *q* are increased in the absence of EAAC1.

**Figure 3.**
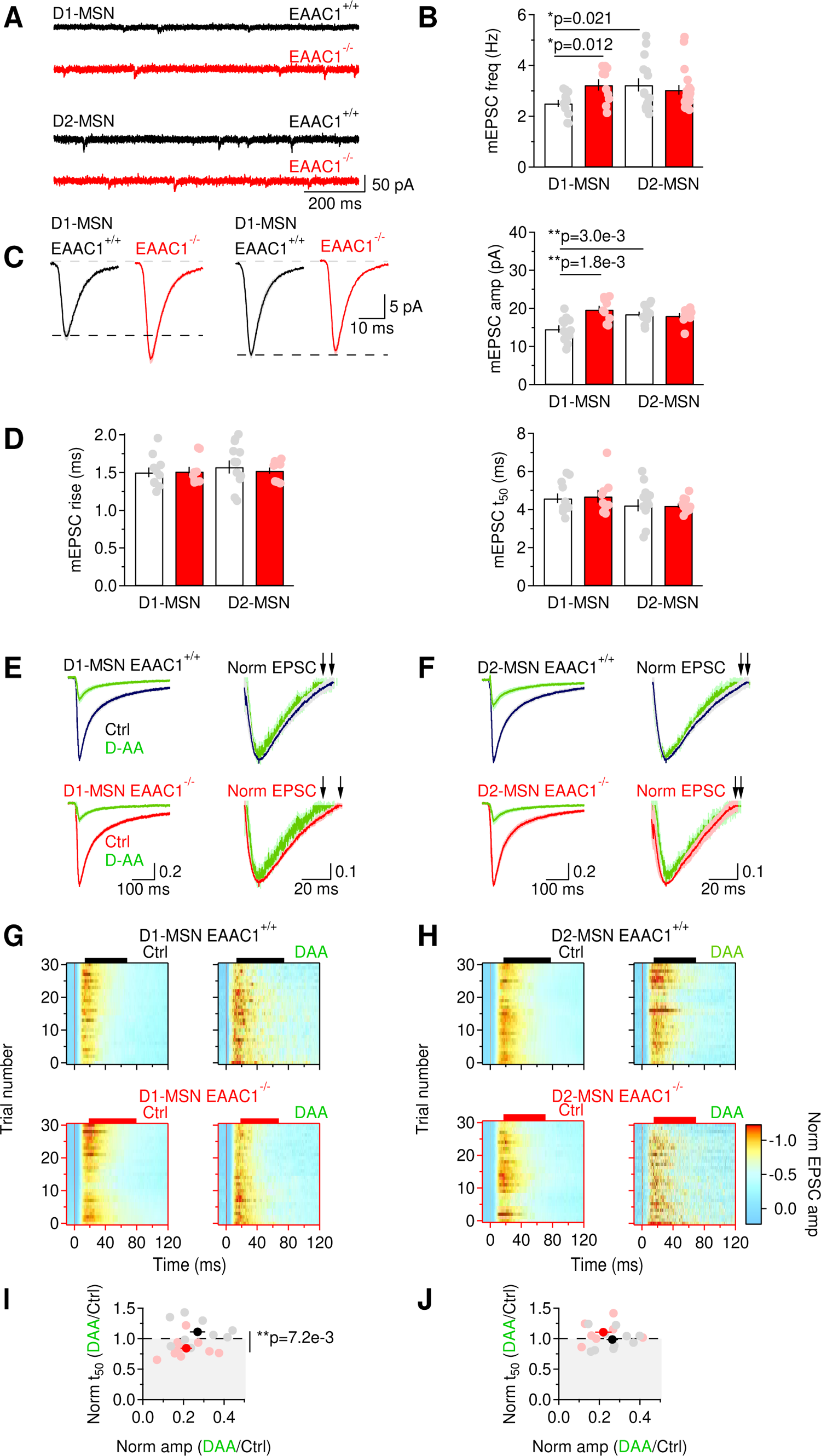
EAAC1 limits excitation in D1-MSNs. **(A)** Sample traces of mEPSCs recordings from DLS MSNs in EAAC1^+/+^ (*black*) and EAAC1^-/-^ mice (*red*), at V_hold_=-70 mV. **(B)** Summary graph of the mEPSC frequency in DLS MSNs. In EAAC1^+/+^ mice, the mEPSC frequency is higher in D2-MSNs (n=12) compared to D1-MSNs (n=15). In EAAC1^-/-^ mice, there is no significant difference in the mEPSC frequency between D1-MSNs (n=10) and D2-MSNs (n=19), but the mEPSC frequency is higher in D1-MSNs compared to EAAC1^+/+^ mice. Data shown in B-D were collected from the same cells. **(C)** *Left,* Average mEPSCs from EAAC1^+/+^ and EAAC1^-/-^ D1- and D2-MSNs. *Right,* Summary of the mEPSC amplitude. In EAAC1^+/+^ mice, the mEPSC amplitude is larger in D2-compared to D1-MSNs. In EAAC1^-/-^ mice, there is no significant difference in the mEPSC amplitude between D1- and D2-MSNs, but the mEPSC amplitude is larger in D1-MSNs compared to EAAC1^+/+^ mice. **(D)** Summary of the mEPSC rise (*left*) and 50% decay time (*right*) measured in D1- and D2-MSNs of EAAC1^+/+^ and EAAC1^-/-^ mice. **(E)** *Left,* NMDA EPSCs from D1-MSNs, recorded in Mg^2+^-free extracellular solution at V_hold_=-70 mV, in control conditions (EAAC1^+/+^ n=9: *black*; EAAC1^-/-^ n=10: *red*) and in the presence of D-AA (100 µM; *green*). *Right,* peak normalized NMDA EPSCs, shown with a y-axis range of 0.5-1. This allows visualizing the time used for the calculations of the t_50_, highlighted by the black arrows. Each trace represents the average of 20 consecutive trials. **(F)** As in E, for D2-MSNs (EAAC1^+/+^ n=10; EAAC1^-/-^ n=7). **(G)** Heat maps showing the time course of the peak normalized NMDA EPSC amplitude in 20 consecutive trials, before (*left*) and after applying D-AA (*right*), in D1-MSNs from EAAC1^+/+^ (*top*) and EAAC1^-/-^ mice (*bottom*). The colored bar at the top of each graph provides a measure of the average t_50_ for each cell. **(H)** As in G, for D2-MSNs. **(I)** Summary scatter plot showing that D-AA reduced the NMDA EPSC amplitude to the same extent in EAAC1^+/+^ and EAAC1^-/-^ D1-MSNs but reduced the t_50_ only in D1-MSNs from EAAC1^-/-^ mice. **(J)** As in I, for D2-MSNs. Data represent mean ± SEM.

The role of EAAC1 in modulating evoked glutamatergic transmission has been previously studied in the hippocampus, where this neuronal transporter has been shown to limit spillover activation of NMDA receptors (Diamond 2001; Scimemi, Tian, and Diamond 2009). Whether this is a general property of EAAC1 and whether EAAC1 exerts similar effects on evoked excitatory transmission in the DLS remains unclear, given that the contribution of NMDA receptors to excitatory synaptic transmission in the DLS is much less pronounced than in the hippocampus (Sung, Choi, and Lovinger 2001). This was confirmed by our own experiments, showing that the NMDA/AMPA ratio in D1-MSNs was 2-4 times smaller than we observed in the hippocampus (McCauley et al. 2020) (**Supp. Fig. 3-1A, B**). If, despite the paucity of NMDA receptors in the DLS, EAAC1 limited their activation by glutamate spillover, we would expect NMDA EPSCs in MSNs of EAAC1^-/-^ mice to decay faster in the presence of the low-affinity, competitive NMDA receptor antagonist D-AA (Diamond 2001). This rationale is based on evidence that: *(i)* D-AA competes with glutamate for binding to NMDA receptors (Clements et al. 1992), and *(ii)* D-AA preferentially blocks receptors activated by small and slow glutamate transients (i.e., peri- and extra-synaptic NMDA receptors) (Diamond 2001). To test this hypothesis, we perfused brain slices with a Mg^2+^-free extracellular solution, and recorded NMDA EPSCs from MSNs voltage clamped at −70 mV, a potential that allows glutamate binding and translocation via post-synaptic EAAC1 (Wadiche et al. 1995; Arriza et al. 1994). To make a meaningful comparison of the effect of D-AA on the NMDA EPSC decay, we used a sub-saturating concentration of D-AA and confirmed that this reduced the NMDA EPSC amplitude to the same extent in EAAC1^+/+^ and EAAC1^-/-^ mice, as this is mostly accounted for by activation of synaptic NMDA (Clements et al. 1992). D-AA (100 µM) reduced the NMDA EPSC amplitude to similar levels in D1-MSNs (EAAC1^+/+^: ***p=5.2e-8; EAAC1^-/-^: ***p=4.6e-10; EAAC1^-/-^ vs. EAAC1^+/+^ p=0.26) and D2-MSNs of EAAC1^+/+^ and EAAC1^-/-^ mice (EAAC1^+/+^: ***p=6.9e-10; EAAC1^-/-^: ***p=1.2e-6; EAAC1^-/-^ vs. EAAC1^+/+^ p=0.38; **Fig. 3E-J**). Under these experimental conditions, D-AA sped the NMDA EPSC only in D1-MSNs of EAAC1^-/-^ mice (norm t_50_ D-AA/Ctrl D1-MSNs EAAC1^+/+^: p=0.15; EAAC1^-/-^: *p=0.013; EAAC1^-/-^ vs. EAAC1^+/+^ **p=7.2e-3; norm t_50_ D-AA/Ctrl D2-MSNs EAAC1^+/+^: p=0.82; EAAC1^-/-^: p=0.18; EAAC1^-/-^ vs. EAAC1^+/+^ p=0.20; **Fig. 3E-J**). These findings indicate that EAAC1 limits spillover activation of NMDA receptors at excitatory synapses onto D1-MSNs, not D2-MSNs, suggesting the existence of a preferential effect of EAAC1 on these cells.

### EAAC1 strengthens inhibition onto D1-MSNs

One interesting property of EAAC1 is that its expression is not limited to glutamatergic synapses but can also be detected in axonal boutons of GABAergic neurons, where it may serve to supply these neurons with a substrate for GABA synthesis and release (Conti et al. 1998; Rothstein et al. 1994; He et al. 2000). This is because even though GABA can be synthesized *de novo* in axonal boutons, part of it can also be synthesized from glutamate recycled from the extracellular space via glutamate transporters (Scimemi 2014). This recycling pathway differs in complexity depending on whether recycling of extracellular glutamate relies on neuronal or glial glutamate transporters (**Fig. 4A, *left***). Whereas glutamate taken up by EAAC1 is converted into GABA via decarboxylation in the presynaptic terminal, glutamate taken up by glial transporters needs first to be converted into glutamine in the astrocyte cytoplasm, which is shuttled to neurons and converted first into glutamate and ultimately into GABA. There are multiple unknowns about these two recycling pathways. For example, it is not known how much GABA release onto MSNs relies on *de novo* synthesis versus recycling via neuronal or glial glutamate transporters. It is also unknown whether EAAC1 contributes differently to action potential-dependent and -independent GABA release. This is important, given the growing body of work indicating that vesicles mediating spontaneous and evoked inhibitory synaptic transmission are partially segregated among synapses and may utilize partially different molecular machineries (Horvath et al. 2020; Sara et al. 2011; Wang et al. 2022).

**Figure 4.**
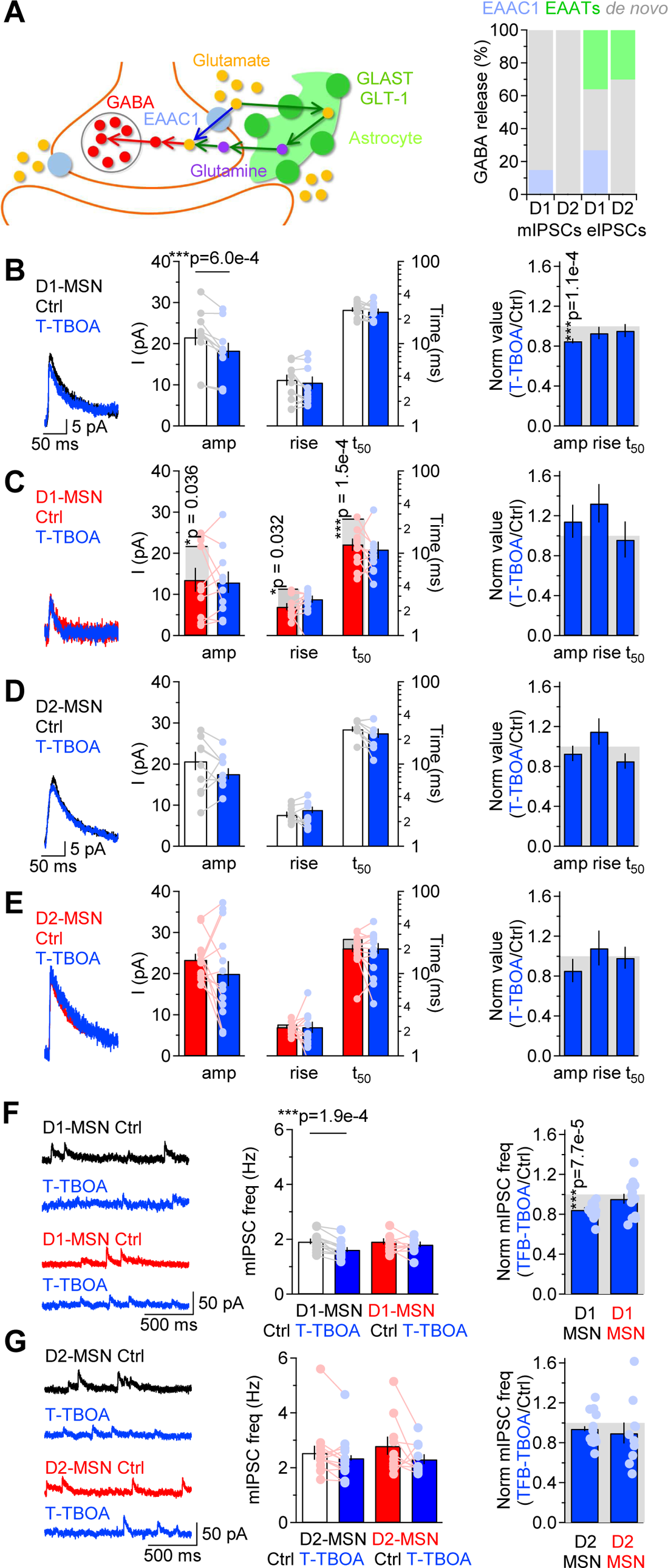
EAAC1 strengthens action potential-independent GABAergic inhibition onto D1-MSNs. **(A)** *Left,* Schematic representation for routes of glutamate uptake via neuronal and glial glutamate transporters. *Right,* Summary of the contribution of glutamate uptake via EAAC1, EAATs and *de novo* GABA synthesis to action potential-independent GABA release. **(B)** *Left,* Example of GABA mIPSCs recorded from D1-MSNs of EAAC1^+/+^ mice in control conditions (*black*) and in the presence of T-TBOA (1 µM; *blue*), at V_hold_=40 mV. *Center,* Summary of the mIPSC amplitude and kinetics recorded before and after T-TBOA (n=11). *Right,* Summary of the relative effect of T-TBOA on the mIPSC amplitude and kinetics. T-TBOA induced a significant reduction in the amplitude of mIPSCs recorded from D1-MSNs. **(C-E)** As in B, for DLS D1-MSNs of EAAC1^-/-^ mice (n=10), and D2-MSNs of EAAC1^+/+^ mice (n=14), and EAAC1^-/-^ mice (n=13), respectively. In panels C, E we also included data collected from EAAC1^+/+^ mice to show that loss of EAAC1 leads to smaller and faster mIPSCs in D1-MSNs, whereas no significant difference is detected between D2-MSNs of EAAC1^+/+^ and EAAC1^-/-^ mice. **(F)** Summary of the effect of T-TBOA on the mIPSC frequency in D1-MSNs of EAAC1^+/+^ and EAAC1^-/-^ mice, with representative traces (*left*), raw data (*center*) and normalized values (*right*). **(G)** As in **F**, for D2-MSNs.

We addressed these concerns first by recording mIPSCs in MSNs from EAAC1^+/+^ and EAAC1^-/-^ mice. The mIPSCs were recorded in the presence of voltage-gated sodium channel, AMPA and NMDA receptor blockers (TTX 1 µM, NBQX 10 µM and APV 50 µM, respectively) from MSNs voltage clamped at 40 mV. By holding cells at this depolarized potential, we limited potential confounding effects due to post-synaptic uptake via EAAC1, which is inhibited by membrane depolarization (Wadiche et al. 1995). We recorded mIPSCs before and after applying T-TBOA (1 µM), a broad-spectrum glutamate transporter antagonist (Tsukada et al. 2005). Our results showed that T-TBOA decreased the mIPSC amplitude in D1-MSNs from EAAC1^+/+^ mice by ∼15%, without changing the mIPSC kinetics (EAAC1^+/+^ D1-MSN mIPSC amplitude, Ctrl vs. T-TBOA: **p=5.9e-4; rise: p=0.32; t_50_: p=0.19). By contrast, the amplitude and kinetics of mIPSCs recorded from D1-MSNs from EAAC1^-/-^ mice were not altered by T-TBOA (EAAC1^-/-^ D1-MSN mIPSC amplitude, Ctrl vs. T-TBOA: p=0.76; rise: p=0.13; t_50_: p=0.55; **Fig. 4B,C**). The mIPSC amplitude and kinetics in D2-MSNs were not altered by T-TBOA, in EAAC1^+/+^ and EAAC1^-/-^ mice (EAAC1^+/+^ D2-MSN mIPSC amplitude, Ctrl vs. T-TBOA: p=0.06; rise: p=0.19; t_50_: p=0.20; EAAC1^-/-^ D2-MSN mIPSC amplitude, Ctrl vs. T-TBOA: p=0.25; rise: p=0.78; t_50_: p=0.72; **Fig. 4D,E**). These data indicate that: *(i)* EAAC1 is the sole glutamate transporter contributing to filling of vesicles used for action potential-independent GABA release onto D1-MSNs; *(ii)* neither EAAC1 nor astrocytic glutamate transporters contribute to filling of vesicles released spontaneously onto D2-MSNs. These conclusions were consistent with two additional observations. *First,* in control conditions, the mIPSC amplitude, rise and decay time were smaller in D1-MSNs from EAAC1^-/-^ compared to EAAC1^+/+^ mice (mIPSC amplitude: *p=0.04; rise: *p=0.03; t_50_: ***p=1.5e-4; **Fig. 4B,C**). By contrast, the amplitude, rise and 50% decay time of mIPSCs recorded from D2-MSNs were similar in EAAC1^+/+^ and EAAC1^-/-^ mice (amplitude: p=0.32; rise: p=0.24; t_50_: p=0.06; **Fig. 4D,E**). *Second*, T-TBOA reduced the mIPSC frequency only in D1-MSNs from EAAC1^+/+^ (***p=1.9e-4), not EAAC1^-/-^ mice (p=0.22; **Fig. 4F**). There was no significant effect of T-TBOA on the mIPSC frequency in D2-MSNs (EAAC1^+/+^: p=0.13; EAAC1^-/-^: p=0.14; **Fig. 4G**). Therefore, EAAC1 shapes not only glutamatergic transmission but also action potential-independent GABA release onto D1-MSNs.

We next analyzed the effect of T-TBOA on evoked IPSCs (**Fig. 5A-D**). T-TBOA reduced the IPSC amplitude in MSNs of EAAC1^+/+^ and EAAC1^-/-^ mice, without altering their rise and 50% decay time. The reduction of the IPSC amplitude induced by T-TBOA was ∼63% in D1-MSNs of EAAC1^+/+^ mice and ∼36% in D1-MSNs of EAAC1^-/-^ mice (*p=0.02; **Fig. 5A, B, E**). These results suggest that EAAC1 contributes to ∼27% of the quantal size of vesicles released in an action potential-dependent manner onto D1-MSNs (i.e., 63%-36%), with a ∼36% contribution by other glial transporters and ∼37% contribution by *de novo* synthesis (i.e., 100%-63%; **Fig. 4A, *right***). In D2-MSNs, T-TBOA reduced the IPSC amplitude by ∼30% in EAAC1^+/+^ and EAAC1^-/-^ mice, suggesting that GABA supply for evoked release onto D2-MSNs relies for ∼70% on *de novo* synthesis and for ∼30% on glial glutamate transporters, with no contribution of EAAC1 (**Fig. 4A, *right,* Fig. 5C-E**). In separate experiments, we confirmed that these results were not biased by a potential presynaptic effect of T-TBOA on release probability at GABAergic synapses, because T-TBOA did not change the IPSC PPR (EAAC1^+/+^ D1-MSN PPR Ctrl vs. T-TBOA: p=0.19; EAAC1^-/-^ D1-MSN PPR Ctrl vs T-TBOA: p=0.73; EAAC1^+/+^ D2-MSN PPR Ctrl vs. T-TBOA: p=0.40; EAAC1^-/-^ D2-MSN PPR Ctrl vs. T-TBOA: p=0.16; **Fig. 5A-D, H**). Together, these findings suggest that EAAC1 enhances spontaneous and evoked GABAergic transmission onto D1-MSNs, without altering GABAergic inhibition onto D2-MSNs (**Fig. 5E-H**).

**Figure 5.**
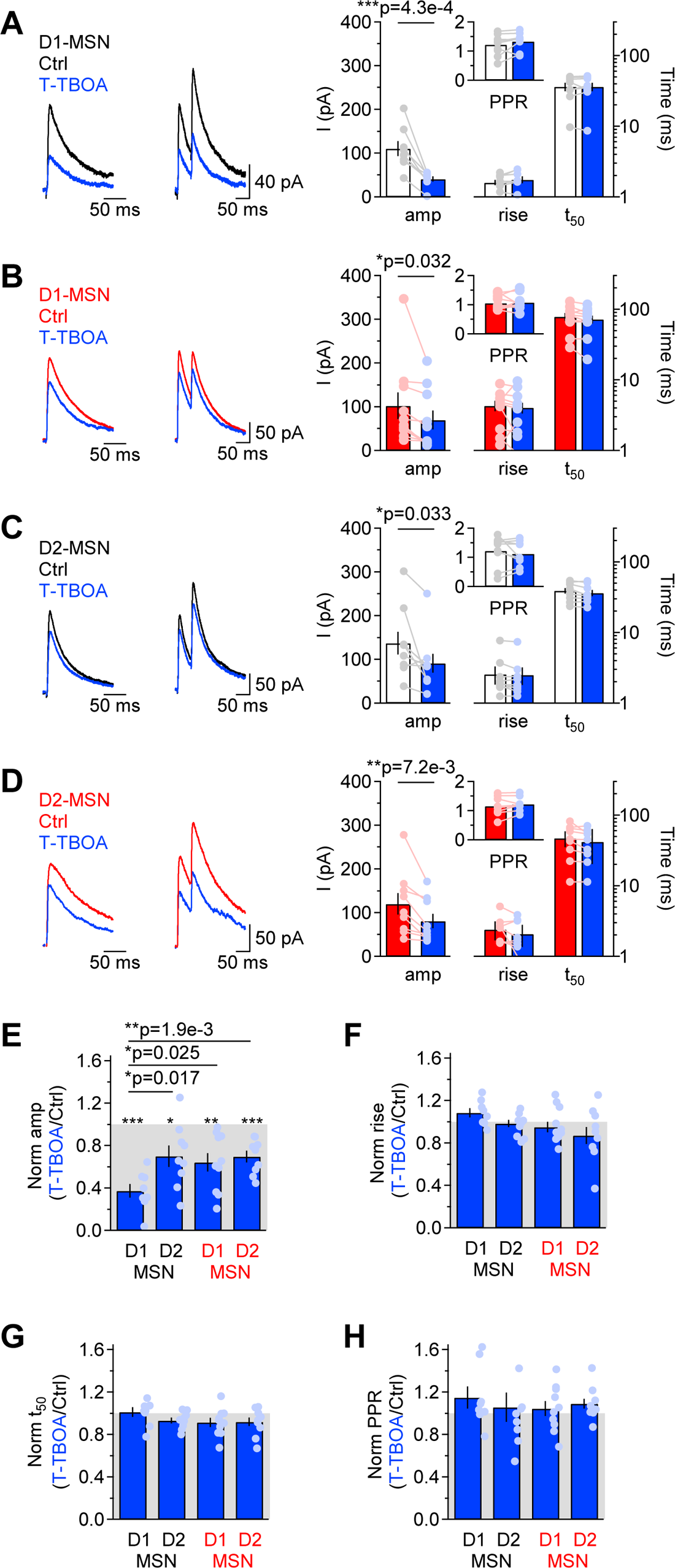
EAAC1 strengthens action potential-dependent GABAergic inhibition onto D1-MSNs. **(A)** *Left,* Example of evoked single and paired IPSCs recorded from D1-MSNs of EAAC1^+/+^ mice in control conditions (*black*) and in the presence of T-TBOA (1 µM; *blue*), at V_hold_=40 mV. *Center,* Summary of the IPSC amplitude and kinetics recorded before and after T-TBOA (n=9). The inset represents the summary values of the PPR. **(B-D)** As in A, for DLS D1-MSNs (n=10) and D2-MSNs of EAAC1^+/+^ mice (n=8), and EAAC1^-/-^ mice (n=10), respectively. **(E-H)** Summary of the relative effect of T-TBOA on the IPSC amplitude and kinetics. T-TBOA induced a significant reduction on the IPSC amplitude, but not of the rise time, decay time and PPR. T-TBOA reduced the IPSC amplitude in D1-MSNs of EAAC1^+/+^ mice more than in any other type of MSNs tested in the experiments. Data represent mean ± SEM.

### EAAC1 strengthens reciprocal inhibition between D1-MSNs

MSNs form extensive collaterals within the DLS. Accordingly, the dendritic field of each MSN extends over a volume that contains >2,800 MSNs, and forms ∼1,200 – 1,800 synaptic contacts onto these cells (Wilson 2007). Functional studies suggest that each D1-MSN has a connection rate of 26% and 6% with other D1- and D2-MSNs, respectively (Taverna, Ilijic, and Surmeier 2008; Tecuapetla et al. 2009). By contrast, each D2-MSN has a connection rate of 28% and 36% with other D1- and D2-MSNs, respectively (Tecuapetla et al. 2009; Taverna, Ilijic, and Surmeier 2008). The presence of such an intricate inhibitory network of connections is thought to be important for regulating the excitability of MSNs and, more generally, the output of the striatum and the execution of coordinated motor behaviors (Burke, Rotstein, and Alvarez 2017; Dobbs et al. 2016). To determine to which extent EAAC1 shapes lateral inhibition among different types of MSNs, we performed bilateral stereotaxic injections of a conditional ChR2-expressing AAV in the DLS of D1^Cre/+^:Ai9^Tg/Tg^ or A2A^Cre/+^:Ai9^Tg/Tg^ mice (**Fig. 6A, *left***). Three weeks later, we prepared acute coronal slices, patched D1- or D2-MSNs, identified for their expression of the tdTomato reporter, and used blue light stimuli for full-field activation of ChR2 expressed in either D1- or D2-MSNs (**Fig. 6A, *right***). We set the intensity of the blue light stimulation to a power of ∼250 µW, measured at the sample plane (**Supp. Fig. 6-1A**). Each stimulation lasted 5 ms, a duration that did not evoke any adaptation in post-synaptic, ChR2-mediated photocurrents (**Supp. Fig. 6-1B**), and was repeated every 30 s, to allow for full recovery of ChR2 from activation (**Supp. Fig. 6-1D**). The ChR2 photocurrents, isolated in the presence of TTX (1 µM), reversed at ∼20 mV (**Supp. Fig. 6-1C**) and their amplitude did not change in the presence of the GABA_A_ receptor antagonist picrotoxin (100 µM). Therefore, we used 20 mV as the holding potential to record optogenetically-evoked IPSCs (oIPSCs), isolated pharmacologically by adding NBQX (10 µM) and APV (50 µM) to the external solution. Under these experimental conditions, the oIPSCs were completely blocked by the GABA_A_ receptor antagonist picrotoxin (100 µM). To confirm the accuracy of the stereotaxic injections in the DLS, at the completion of the recordings, we fixed the slices and imaged them using a confocal microscope (**Fig. 6B**). Our experiments showed that the oIPSC amplitude was smaller at D1-D1 synapses from EAAC1^-/-^ compared to EAAC1^+/+^ mice, suggesting that EAAC1 might enhance synaptic inhibition at these synapses (*p=0.04; **Fig. 6D, H**). By contrast, the amplitude of oIPSCs evoked at D2-D1 synapses was similar across the two genotypes (p=0.65; **Fig. 6E, I**). The oIPSC amplitude was also similar at D1-D2 (p=0.77; **Fig. 6F, J**) and D2-D2 synapses of EAAC1^+/+^ and EAAC1^-/-^ mice (p=0.51; **Fig. 6G, K**). Although T-TBOA (1 µM) reduced the oIPSC amplitude at all collateral MSN synapses, its effect was largest at D1-D1 synapses (**Fig. 6D-K**). Accordingly, the reduction of the oIPSC amplitude induced by T-TBOA at D1-D1 synapses was ∼53% in EAAC1^+/+^ mice and ∼29% in EAAC1^-/-^ mice (**p=9.1e-3; **Fig. 6H**). This suggests that GABAergic inhibition between D1-MSNs relies for ∼24% on glutamate uptake via EAAC1 (i.e., 53%-29%), and for ∼29% on glutamate uptake mediated by glial transporters (**Fig. 6C**). The reduction of the oIPSC amplitude induced by T-TBOA was similar at D2-D1 (p=0.72), D1-D2 (p=0.26) and D2-D2 synapses in EAAC1^+/+^ and EAAC1^-/-^ mice (p=0.86; **Fig. 6D-K**). These findings suggest that glial glutamate transporters contribute to 33-44% of GABA release across different types of collateral synapses formed between MSNs (**Fig. 6C**).

**Figure 6.**
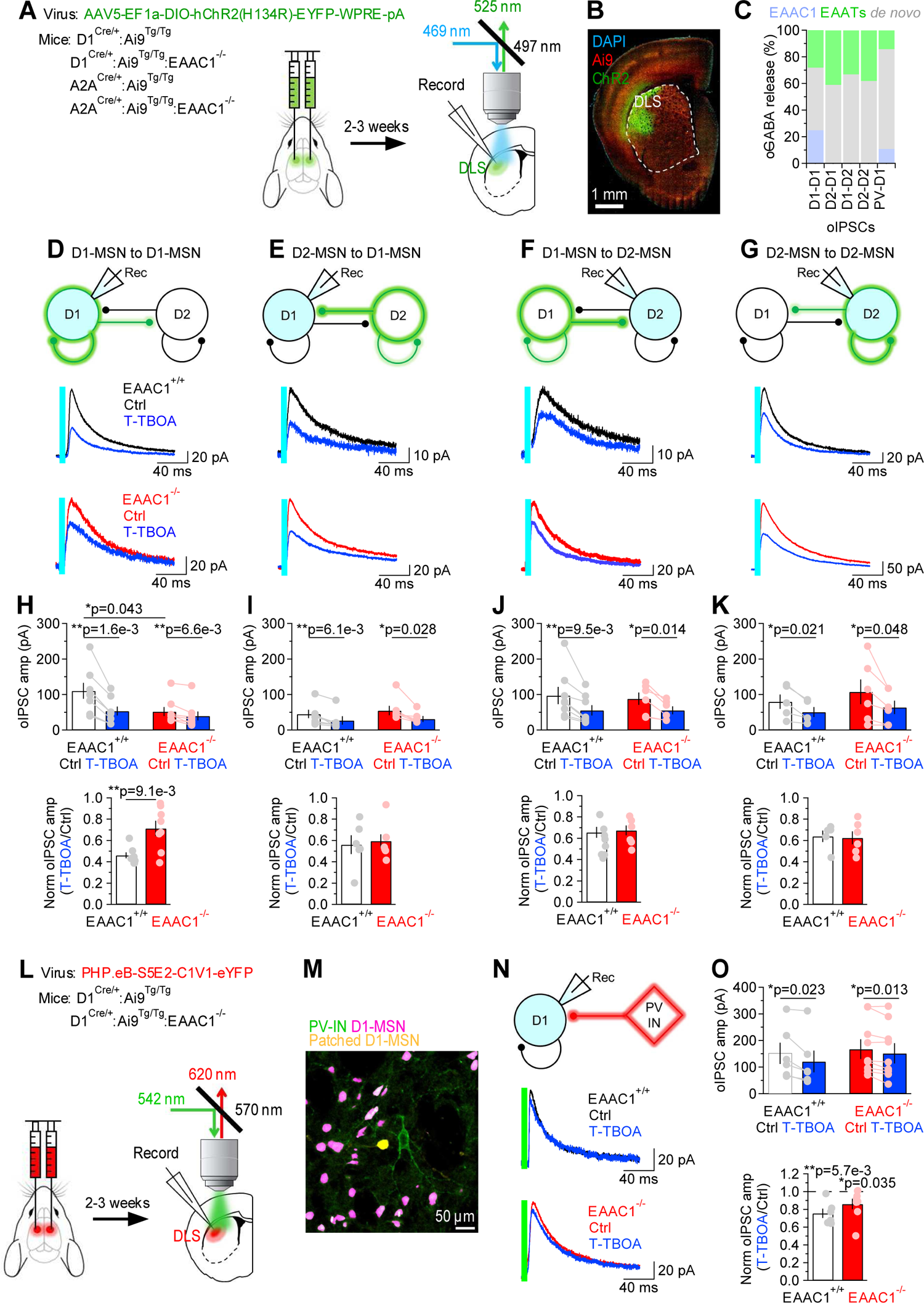
EAAC1 strengthens GABAergic inhibition at D1-D1 synapses. **(A)** Schematics of viral transduction in the DLS and light evoked stimulation of ChR2-transfected MSNs in slices. **(B)** Confocal image of mouse coronal slice transfected with ChR2. **(C)** *Right,* Summary of the contribution of glutamate uptake via EAAC1, EAATs and *de novo* GABA synthesis to action potential-dependent GABA release evoked by light activation of ChR2 at different sets of synapses. **(C)** Summary of the contribution of different types of glutamate transporters and *de novo* GABA synthesis to GABA released during oIPSCs. **(D)** *Top,* Schematic representation of the experimental design. The patched cell was voltage clamped at V_hold_=20 mV, the reversal potential of ChR2 photocurrents. *Middle,* Example of oIPSCs recorded from D1-MSNs in response to optogenetic stimulation of GABA release from other D1-MSNs, in mice expressing EAAC1. Each trace represents the average of 20 consecutive trials. *Bottom,* As in the middle panel, for EAAC1^-/-^ mice. **(E-G)** As in D, for D2-D1 (E), D1-D2 (F) and D2-D2 synapses (G). **(H)** *Top,* In-cell comparison of D1-D1 oIPSCs in EAAC1^+/+^ (n=9) and EAAC1^-/-^ mice (n=6), before and after T-TBOA (1 µM; *blue*). *Bottom,* Summary of the peak normalized oIPSC amplitude. **(I)** As in H, for D2-D1 oIPSCs in EAAC1^+/+^ (n=7) and EAAC1^-/-^ mice (n=6). **(J)** As in H, for D1-D2 oIPSCs in EAAC1^+/+^ (n=8) and EAAC1^-/-^ mice (n=6). **(K)** As in H, for D2-D2 oIPSCs in EAAC1^+/+^ (n=5) and EAAC1^-/-^ mice (n=6). **(L)** Schematics of viral transduction in the DLS and light evoked stimulation of C1V1-transfected PV-INs in slices. **(M)** Confocal image of tdTomato expressing D1-MSNs (magenta) and C1V1 transfected PV-INs (green) in a slice from which we recorded oIPSCs from a D1-MSN (yellow). **(N)** As in D, for oIPSCs recorded from D1-MSNs in response to green light activation of C1V1 expressed in PV-INs. **(O)** As in H, for oIPSCs recorded in response to C1V1 activation of GABA release from PV-INs to D1-MSNs. Data represent mean ± SEM.

Although the FISH data suggest that EAAC1 is mostly expressed in MSNs (**Fig. 1**), we asked whether it could also shape GABAergic inhibition onto D1-MSNs originating from other cell types. In addition to receiving reciprocal inhibition from D1/2-MSNs, D1-MSNs receive strong feedforward inhibition from a class of interneurons, the fast-spiking parvalbumin interneurons (PV-INs), which *In vivo* can fire action potentials at high frequency (Burke, Rotstein, and Alvarez 2017; Tepper, Koos, and Wilson 2004; Tepper et al. 2010; Tepper, Wilson, and Koos 2008). To determine whether EAAC1 also altered GABAergic inhibition from PV-INs onto D1-MSNs, we performed whole-cell patch clamp recordings from tdTomato-expressing D1-MSNs in mice that received stereotaxic injections of a viral construct that allowed expression of the red-shifted opsin C1V1 in PV-INs (PHP.eB-S5E2-C1V1-eYFP; **Fig. 6L, M**) (Vormstein-Schneider et al. 2020). In these experiments, T-TBOA reduced the oIPSC amplitude similarly in EAAC1^+/+^ and EAAC1^-/-^ mice (**p=5.7e-3 and *p=0.035, respectively; EAAC1^+/+^ vs. EAAC1^-/-^: p=0.15; **Fig. 6N, O**). This finding suggests that EAAC1 does not contribute significantly to GABA release at PV to D1-MSN synapses (**Fig. 6C**).

Together, these results indicate that glial glutamate transporters enhance inhibition at homo- and hetero-synaptic contacts formed between different types of MSNs and, to a lesser extent, at synapses formed between PV-INs and D1-MSNs (**Fig. 6C**). By contrast, EAAC1 significantly enhances synaptic inhibition only at homosynaptic GABAergic contacts formed between D1-MSNs.

### EAAC1 limits the firing output of D1-MSNs

An essential aspect of information processing is the ability to transform synaptic inputs into action potential outputs, allowing the DLS to control the activity of its target regions. The data obtained so far provide an opportunity to shed light on the functional role of homosynaptic, reciprocal inhibition between D1-MSNs. For this reason, we asked how the changes in synaptic excitation and reciprocal inhibition between D1-MSNs induced by EAAC1 shape the firing output of these cells. To answer this, we used the NEURON platform and performed modeling experiments on reconstructed D1-MSNs from EAAC1^+/+^ and EAAC1^-/-^ mice, to account for the different morphological features of these cells (**Fig. 2**) (Hines and Carnevale 1997). In these simulations, the passive and active membrane properties of D1-MSNs were consistent with those of D1-MSNs in EAAC1^+/+^ and EAAC1^-/-^ mice (**Supp. Fig. 7-1**), and the weight of each excitatory and inhibitory input was set to be consistent with our experimental data (**Supp. Fig. 7-2, 7-3, 7-4**). We allowed each D1-MSN to receive 100 excitatory and 100 inhibitory inputs, randomly distributed along the dendrites, activated at a range of frequencies consistent with the range of synaptic inputs received by the striatum *in vivo*. We analyzed the effect of these parameter manipulations on firing rates of D1-MSNs across the theta (*∼*8 Hz) and beta range (*∼*20 Hz), also representative of the firing activity detected in the striatum *in vivo* (**Fig. 7A, B**) (Berke et al. 2004; Pennartz et al. 2009). We then asked whether and how EAAC1 altered the offset and the gain in the input-output relationship of D1-MSNs (**Fig. 7C, *left and center***). Briefly, changing the offset allows D1-MSNs to subtract basal levels of synaptic activity, altering the range of input stimulation frequencies that evoke firing (an additive/subtractive operation). Changing the gain alters the sensitivity of D1-MSNs to varying levels of synaptic input rates, while preserving the range over which these inputs evoke spiking activity (a multiplicative/divisive operation) (Mitchell and Silver 2003). In the absence of synaptic inhibition, EAAC1 induced a modest increase in offset, which became more pronounced as the inhibition rate increased (**Fig. 7D**). With increasing rates of inhibition onto D1-MSNs, EAAC1 also decreased the gain, and therefore switched from having a purely additive effect to also having a divisive effect (**Fig. 7D, *right***). Conversely, when excitation was low, EAAC1 reduced the basal firing rate of D1-MSNs, with a negligible effect on gain (**Fig. 7E, *left***). As excitation increased, the increase in gain becomes more evident and is associated with a slight decrease in the basal firing rate of D1-MSNs (**Fig. 7E, *right***). The overall effect of EAAC1 across a broader range of activity of E/I is to introduce a frequency-dependent increase in offset and a frequency-independent decrease in gain (**Fig. 7C, *right***). Together, these results suggest that EAAC1 can perform both subtractive and divisive operations, depending on the rate of incoming E/I onto D1-MSNs. The higher is the rate of synaptic E/I onto D1-MSNs, the greater is the reduction of the firing output of D1-MSNs by EAAC1. By modulating E/I, EAAC1 narrows the dynamic range and reduces the sensitivity of D1-MSNs to incoming inputs.

**Figure 7.**
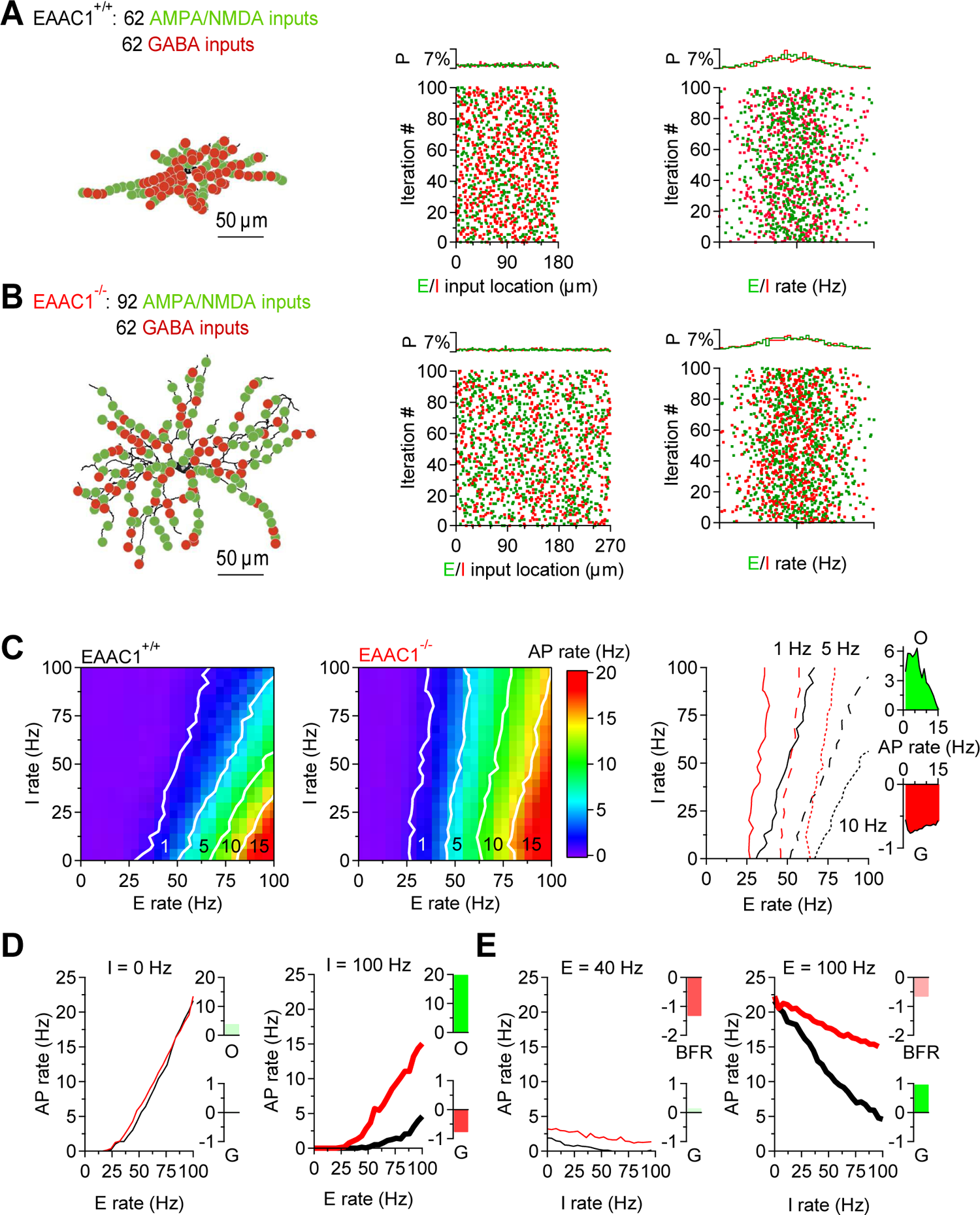
EAAC1 reduces the firing output of D1-MSNs in a realistic compartmental model of D1-MSNs. **(A)** *Left,* Representation of a biocytin-filled D1-MSN in EAAC1^+/+^ mice, with excitatory (*green*) and inhibitory inputs (*red*) randomly distributed along all dendrites. *Center,* Spatial distribution of excitatory and inhibitory inputs along the length of the dendrites, *Right,* Schematic representation of the instantaneous frequency of activation of excitatory and inhibitory inputs in each of the 100 simulation iterations. The activation frequency for these inputs was set to have a Gaussian distribution. **(B)** As in A, for a biocytin-filled D1-MSN in EAAC1^-/-^ mice. **(C)** *Left,* Heat map showing the firing output of D1-MSNs in EAAC1^+/+^ mice for different combinations of activity for excitatory (*x-axis*) and inhibitory inputs (*y-axis*). *Center*, as in the right panel, for EAAC1^-/-^ mice. *Right,* Overlaid contour lines for action potential firing at 1, 5 and 10 Hz in D1-MSNs from EAAC1^+/+^ and EAAC1^-/-^ mice. **(D)** *Left,* Output firing frequency-input frequency relationships for synaptic excitation for D1-MSNs of EAAC1^+/+^ (*black*) and EAAC1^-/-^ mice (*red*), in the absence (*thin curves*) of synaptic inhibition. *Right,* As in the left panel for high inhibition levels (i.e., 100 Hz; *thick curves*). **(E)** As in D, to compare the effect of inhibition at varying levels of excitation in D1-MSNs of EAAC1^+/+^ and EAAC1^-/-^ mice. Data represent means from 100 simulations.

### The role of reciprocal inhibition between D1-MSNs in reward-based behaviors

The activity of MSNs is important for the execution of coordinated movements and reward-based behaviors (Freeze et al. 2013; Kravitz et al. 2010; Lobo et al. 2010; Kravitz, Tye, and Kreitzer 2012). Although heterosynaptic inhibition from D2-MSNs has been shown to inhibit action potential firing in D1-MSNs (Dobbs et al. 2016), much less is known about the functional significance of homosynaptic inhibition between D1-MSNs. Computational models suggest that this form of inhibition might be involved in driving coherence or synchronizing functional units of information processing in the striatum, known as ensembles (Humphries and Prescott 2010; Humphries, Wood, and Gurney 2010, 2009; Moyer et al. 2014; Ponzi and Wickens 2012, 2010; Ponzi and Wickens 2013; Yim, Aertsen, and Kumar 2011). If the striatum truly operates as a collection of ensembles driving specific behaviors, one may hypothesize that the reduction in synaptic excitation and reciprocal inhibition among D1-MSNs might increase the propensity of mice to switch between different reward-based behaviors. We tested this hypothesis using a simple probabilistic reward lever press task, in which we trained mice to receive a water reward at different probability (P_rew_=0.5‖0.9; **Fig. 8A-E**). In the training sessions, in both EAAC1^+/+^ and EAAC1^-/-^ mice, the number of collected rewards was proportional to the reward probability, and inversely related to the number of level presses (**Fig. 8B, D**). No statistically significant difference between the EAAC1^+/+^ and EAAC1^-/-^ mice was detected (**Fig. 8C, E**). We then ran test sessions where the reward probability was changed every 5-75 s (**Fig. 8F-N**). In these experiments, we still detected an inverse relationship between the number of rewards/lever presses and the reward probability (**Fig. 8F-I**). When the switch time was short (<15 s), EAAC1^+/+^ and EAAC1^-/-^ mice collected a similar number of rewards (**Fig. 8K, L**) and performed a similar number of lever presses (**Fig. 8M, N**) and. As the switch time increased (30-75 s), EAAC1^-/-^ mice outperformed EAAC1^+/+^ mice, collecting more rewards at low and high reward probabilities (**Fig. 8L, N**). Together, these findings suggest that increased excitation onto D1-MSNs and reciprocal inhibition among D1-MSNs limit rapid execution of reward-based behaviors and action switching.

**Figure 8.**
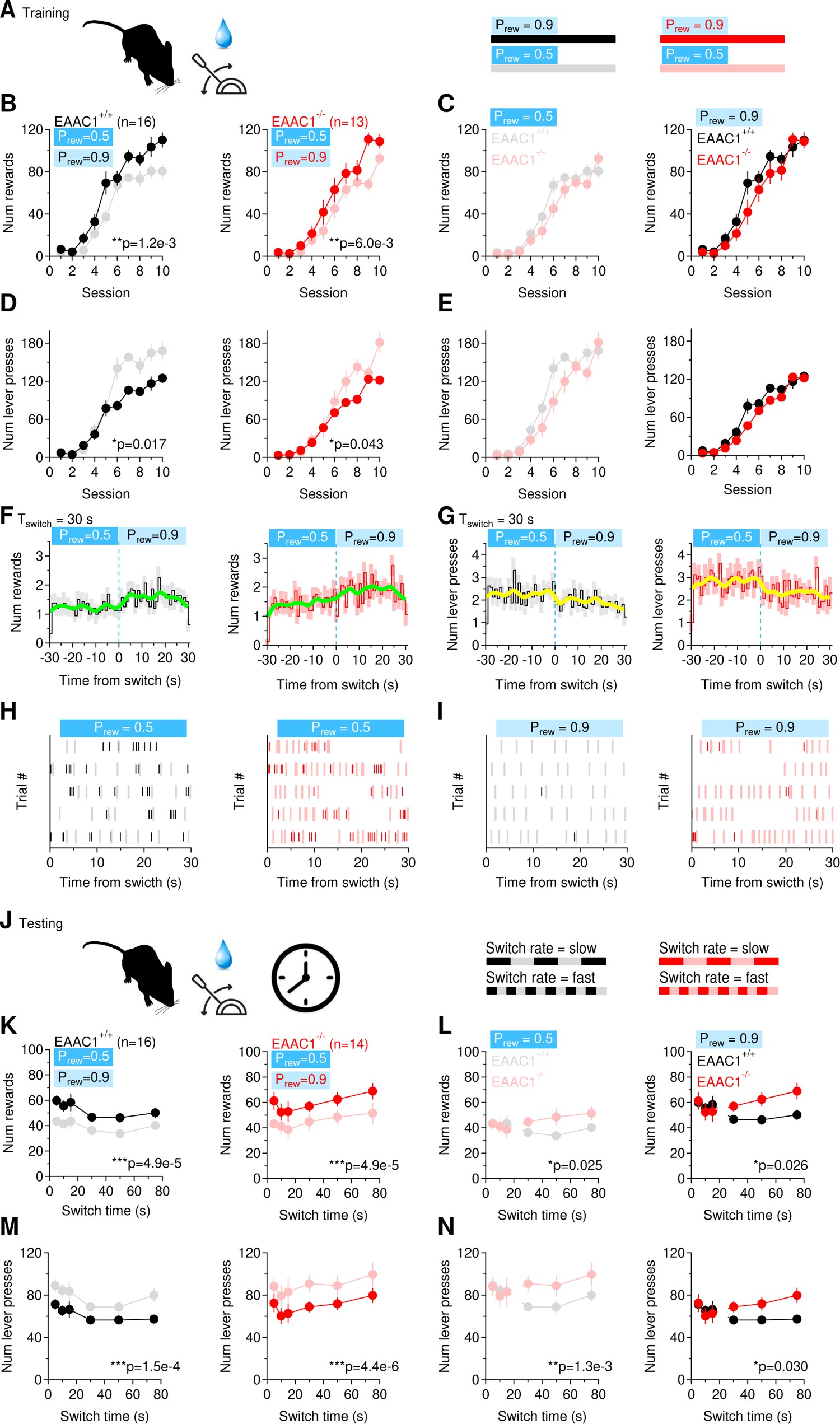
EAAC1 shapes fast-switching behaviors. **(A)** Schematic representation of the training sessions, with water rewards delivered at two different reward probabilities (P_rew_), during each 5 min session. **(B)** Number of rewards collected by EAAC1^+/+^ (n=16) and EAAC1^-/-^ mice (n=16), at P_rew_=0.5 and P_rew_=0.9 over the course of 10 training sessions. **(C)** Comparison between EAAC1^+/+^ and EAAC1^-/-^ mice for the data shown in B. **(D, E)** As in B, C, showing the total number of lever presses performed by the mice. **(F)** Temporal distribution of the number of rewards collected by EAAC1^+/+^ (*left*) an EAAC1^-/-^ mice (*right*) when the reward probability was switched from P_rew_=0.5 to P_rew_=0.9. The thick green line represents a binomial smoothing of the mean. **(G)** As in F, for the number of lever presses. **(H)** Raster plot showing the temporal distribution of lever presses in a 5-min long trial, in which the reward probability was switched from P_rew_=0.5 to P_rew_=0.9 every 30 s. The raster plots in this panel were collected at P_rew_=0.5. Black/red ticks represent the times when the lever was pressed. Gray/pink ticks represent the times when the water rewards were collected. **(I)** As in H, for P_rew_=0.9. **(J)** Schematic representation of the test session, with water rewards delivered at two different values of P_rew_, switching at different time intervals (5-75 s). **(K-N)** As in B-E, for the test session. Data represent mean ± SEM.

## Discussion

The main finding in this work is that the neuronal glutamate transporter EAAC1 exerts a cell-preferential control of E/I in the DLS: it limits excitation onto D1-MSNs and homosynaptic lateral inhibition between D1-MSNs. By doing so, EAAC1 increases the offset and decreases the gain of the firing output of D1-MSNs. These effects are associated with reduced ability to engage in rapidly switching reward-based behaviors.

### The role of EAAC1 in controlling synaptic integration in D1-MSNs

The DLS, one of the four main subdivisions of the striatum, integrates synaptic inputs from multiple brain regions to control the execution of habitual behaviors (Hunnicutt et al. 2016; Wall et al. 2013). In mice, ∼1/3 of excitatory inputs to the DLS originates from the thalamus (Huerta-Ocampo, Mena-Segovia, and Bolam 2014), and the remaining 2/3 from the frontal associative and sensorimotor cortex (Hunnicutt et al. 2016; Guo et al. 2015; Wall et al. 2013; Pan, Mao, and Dudman 2010). All these inputs provide excitation onto both D1- and D2-MSNs, but sensory inputs preferentially innervate D1-MSNs, whereas motor inputs preferentially target D2-MSNs (Wall et al. 2013). Our findings do not distinguish whether EAAC1 limits excitation at some or all these glutamatergic projection types to the DLS, but multiple lines of evidence indicate that only excitation onto D1-MSNs is affected by EAAC1. *First,* there is a reduced spillover-activation of NMDA receptors in these cells (**Fig. 3**). *Second,* EAAC1 expression is associated with a reduced spine size, quantal size (*q*) and quantal content (*N*) in D1-MSNs (**Fig. 2, 3**). *Third,* the dendritic arbor of D1-MSNs is smaller when EAAC1 is expressed (**Fig. 2**). Since D1-MSNs preferentially process sensorimotor information, these findings suggest that EAAC1 is primarily involved with sensorimotor input integration in the DLS.

How excitation onto D1-MSNs is processed, relayed to other brain regions and ultimately converted into different behavioral outputs depends on multiple factors, including: *(i)* the spatial distribution and activation time of these thalamo-cortical excitatory inputs; *(ii)* the local connectivity and pattern of activity of inhibitory connections formed by striatal interneurons and other MSNs; *(iii)* the intrinsic excitability properties of these cells, which however do not change in the absence of EAAC1 (**Supp. Fig. 7-1**) (Burke, Rotstein, and Alvarez 2017; Preston, Bishop, and Kitai 1980; Somogyi, Bolam, and Smith 1981; Wilson and Groves 1980; Kawaguchi, Wilson, and Emson 1990; Yung et al. 1996; Park, Lighthall, and Kitai 1980). MSNs receive strong peri-somatic feedforward inhibition from fast spiking PV-INs, which is not altered by EAAC1 (Burke, Rotstein, and Alvarez 2017). A large population of 1,200-1,800 inhibitory inputs, however, targets the distal dendrites of these cells, and is formed by axon collaterals of other MSNs. Although these distal inputs are thought to be at a positional disadvantage and weaker than peri-somatic inhibitory inputs, they can change local synaptic integration and the firing output of MSNs (Tunstall et al. 2002; Czubayko and Plenz 2002; Dobbs et al. 2016; Koos and Tepper 1999; Wilson 2007). Accordingly, simultaneous activation of D2-MSNs suppresses D1-MSN firing (Dobbs et al. 2016). Given that the coordinated activity of MSNs accounts for the complementary roles of these cells in the temporal control of movement execution (Kravitz et al. 2010; Cui et al. 2013), an increased spiking of D1-MSNs caused by altered E/I in the absence of EAAC1 could provide an important mechanism to disrupt the coordinated recruitment of MSNs during sensorimotor integration and stereotyped movement execution. This hypothesis is supported by our own previous work, showing that EAAC1^-/-^ mice display an increased grooming frequency (Bellini et al. 2018).

### Physiological implications of offset- and gain-modulation by EAAC1

Understanding the input-output transformations of D1-MSNs is a key step in tying together the effect of EAAC1 on E/I. In the input-output relationships, changes in offset or gain exert different effects on information processing (Abbott et al. 1997; Salinas and Thier 2000; Schwartz and Simoncelli 2001; Prescott and De Koninck 2003; Mehaffey et al. 2005). Changes in offset alter the input detection threshold (Pavlov et al. 2009). Changes in gain alter the sensitivity of a neuron to changes in excitatory inputs, and the range of inputs that can be discriminated (Pavlov et al. 2009). Excitatory afferents to the striatum generate action potentials at different rates. In the absence of inhibitory inputs, EAAC1 increases the offset of D1-MSNs (i.e., it increases their detection threshold), but preserves the gain of their input-output relationship. As the rate of D1-D1 inhibition increases, EAAC1 increases the offset even further, while also reducing the gain (i.e., the sensitivity to changes in excitation rates). That is to say that the input detection threshold of D1-MSNs for thalamo-cortical excitatory inputs is increased and the gain is slightly reduced at increasing levels of reciprocal inhibition among these cells, when EAAC1 is expressed. This flexible offset- and gain-control mechanism, which varies with the frequency of E/I and limits D1-MSNs firing in response to ongoing thalamo-cortical excitation, is lost in EAAC1^-/-^ mice.

Multiple studies suggest that MSNs in the DLS are organized into ensembles that can control the execution of different behaviors through their coordinated activation (Adler et al. 2012; Carrillo-Reid et al. 2011; Carrillo-Reid et al. 2008; Barbera et al. 2016). One of the conceptual models for the mechanisms of action of these functional units posits that D1-MSNs in units driving a specific behavior and D2-MSNs in units inhibiting competing behaviors are simultaneously active (Mink 1996). Reciprocal inhibition between D1-MSNs would then limit the execution of competing actions, whereas reciprocal inhibition between D2-MSNs suppresses the inhibition of the desired behavior (Mink 1996). According to this working model, disrupting reciprocal inhibition between D1-MSNs, as it happens in EAAC1^-/-^ mice, would promote unsynchronized activity across different functional units (Humphries, Wood, and Gurney 2010; Moyer et al. 2014; Ponzi and Wickens 2012, 2010; Ponzi and Wickens 2013; Yim, Aertsen, and Kumar 2011). Would this cause the DLS to remain in a given state and promote the sustained execution of a given behavior, or perhaps allow faster switching between different behaviors? Our experiments support a model in which reciprocal inhibition among D1-MSNs limits action switching, and likely promotes coordinated movement execution (**Fig. 8**). Consistent with this hypothesis, loss-of-function mutations of EAAC1 have been implicated with a neuropsychiatric disease characterized by loss of movement coordination and impulsivity, like OCD (Porton et al. 2013). Therefore, by modulating E/I onto D1-MSNs, EAAC1 may be a key regulator of the activity of functional units in the striatum also implicated with this disease. These mechanisms are particularly important in the context of OCD, as they might contribute to increased impulsivity and hypersensitivity to triggers, respectively (Grassi et al. 2015; Boisseau et al. 2012; Bari and Robbins 2013; Benatti et al. 2014; Ettelt et al. 2007).

### Functional implications of EAAC1 in different brain regions and in the context of OCD

Increasing evidence implicates EAAC1 in the modulation of synaptic transmission in at least two regions of the brain: the hippocampus and DLS (Bellini et al. 2018; Scimemi, Tian, and Diamond 2009; Diamond 2001; Mathews and Diamond 2003). There are commonalities in the mechanisms through which EAAC1 controls synaptic transmission in these two brain regions, because in both EAAC1 acts to limit glutamate escape at excitatory synapses and increases GABA release. However, the functional consequences of these effects can vary, due to existing differences in the molecular landscape of excitatory and inhibitory synapses across brain regions. In CA1 pyramidal cells of the mouse hippocampus, EAAC1 limits glutamate escape onto NMDA receptors and promotes long-term plasticity (Diamond 2001; Scimemi, Tian, and Diamond 2009). In the DLS, EAAC1 also limits glutamate escape onto NMDA receptors. In addition, by preventing activation of mGluRI receptors, it promotes D1 dopamine receptor expression and long-term plasticity (Bellini et al. 2018). EAAC1 contributes to GABA synthesis and release from *stratum oriens* interneurons to CA1 pyramidal cells (Mathews and Diamond 2003), and only at a subset of inhibitory synapses mediating lateral inhibition across D1-MSNs in the striatum. The hippocampus has not been the focus of OCD investigations, but there are indications that structural changes in the hippocampus of patients OCD exist (Reess et al. 2018). Although their implications remain unclear, these findings support emerging evidence that limbic structures like the hippocampus contribute to the pathology of OCD (Milad and Rauch 2012; Menzies, Chamberlain, et al. 2008; Wood and Ahmari 2015; Ullrich et al. 2018). Therefore, all these effects are relevant in the context of OCD, as they may contribute to the heterogeneity of symptoms associated with this disease.

Obviously, it is important to remember that although animal studies provide strong evidence for dysfunctions in glutamatergic transmission at striatal thalamo-cortical synapses in the context of OCD, several other genes in the glutamatergic system, in addition to *SLC1A1*, are associated with the disease (Welch et al. 2007; Shmelkov et al. 2010). Other candidate genes belong to the serotonergic and dopaminergic systems. Furthermore, non-genetic, environmental risk factors are also crucial for the manifestation of OCD, making it a complex polygenic and multifactorial disease (Pauls et al. 2014). Despite this apparently daunting scenario, there is considerable agreement on the implication of cortico-striatal hyperactivity in OCD, suggesting the existence of a possible convergence of different genetic and epigenetic factors in the control of striatal function. Therefore, the findings described here may provide an example of circuit and synaptic dysfunctions shared across different genetic models of OCD.

## Materials and Methods

### Ethics Statement

All experimental procedures were performed in accordance with protocols approved by the Institutional Animal Care and Use Committee at the State University of New York (SUNY) Albany and guidelines described in the National Institutes of Health’s Guide for the Care and Use of Laboratory Animals.

### Mice and genotyping

All mice were group housed and kept under a 12 h light cycle (7:00 AM on, 7:00 PM off) with food and water available *ad libitum*. Constitutive EAAC1 knockout mice (EAAC1^-/-^) were obtained by targeted disruption of the *Slc1a1* gene via insertion of a pgk neomycin resistance cassette in exon 1, as originally described by (Peghini, Janzen, and Stoffel 1997). EAAC1^-/-^ breeders were generated after backcrossing EAAC1^-/-^ mice with C57BL/6 mice for >10 generations, as described by Scimemi et al. (2009). C57BL/6 wild type (WT) and EAAC1^-/-^ mice were identified by PCR analysis of genomic DNA. D1^Cre/+^ mice (RRID: MMRRC_030778-UCD; Stock Tg(Drd1-cre)EY217Gsat/Mmucd) and A2A^Cre/+^ mice (RRID: MMRRC_036158-UCD; Stock B6.FVB(Cg)-Tg(Adora2a-cre)KG139Gsat/Mmucd) (Gong et al. 2007; Gong et al. 2003) were kindly provided by Dr. Gerfen (NIH/NIDDK). In these mice, the protein Cre-recombinase is expressed under the control of the promoter for D1 receptors and the adenosine receptor 2 (A2A), respectively. Neurons expressing A2A receptors also express a high density of D2 receptors, and these two receptors establish reciprocal antagonistic interactions in MSNs (Higley and Sabatini 2010). Ai9^Tg/Tg^ conditional reporter mice (RRID: IMSR_JAX:007909; Stock B6.Cg-Gt(ROSA)26Sortm9(CAG-tdTomato)Hze; Madisen et al., 2010) were kindly provided by Dr. P.E. Forni (SUNY Albany). D1^tdT/+^ mice were purchased from The Jackson Laboratory (RRID: IMSR_JAX:016204; Stock Tg(Drd1a-tdTomato)6Calak). For simplicity, we refer to D1^Cre/+^:Ai9^Tg/Tg^, A2A^Cre/+^:Ai9^Tg/Tg^ and D1^tdT/+^ mice as EAAC1^+/+^ mice. Genotyping was performed on toe tissue samples of P7-10 mice. Briefly, tissue samples were digested at 55°C overnight with shaking at 330 rpm in a lysis buffer containing the following (in mM): 100 Tris base pH 8, 5 EDTA, and 200 NaCl, along with 0.2% SDS and 50 µg/ml proteinase K. Following heat inactivation of proteinase K at 97°C for 10 min, DNA samples were diluted 1:1 with nuclease-free water. The PCR primers used for EAAC1, D1^Cre/+^, A2A^Cre/+^, D1^tdT/+^, and Ai9 genotyping were purchased from Thermo Fisher Scientific, and their nucleotide sequences are listed in **Table 1**. PCR was carried out using the KAPA HiFi Hot Start Ready Mix PCR Kit (Cat# KK2602, KAPA Biosystems, Wilmington, MA). Briefly, 12.5 µl of 2X KAPA HiFi Hot Start Ready Mix was added to 11.5 µl of a diluted primer mix (0.5-0.75 µM final for each primer) and 1 µl of diluted DNA. The PCR cycling protocol for all mutants is described in **Table 2**.

**Table 1.**
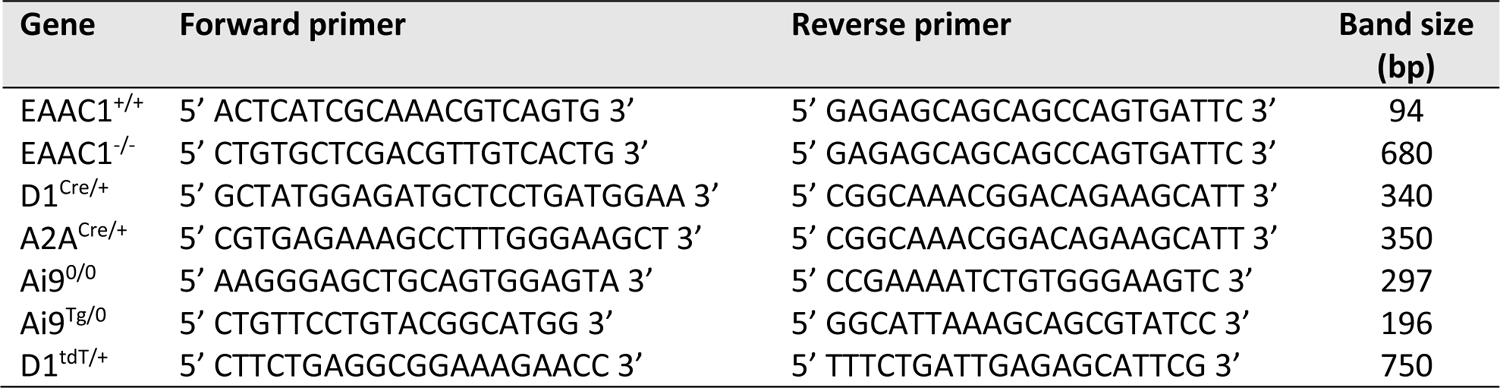
Sequence of genotyping primers

**Table 2.** PCR protocol for all mice

|  | Initiation/<br>melting | Denaturation | Annealing | Elongation | Amplification | Hold |
| --- | --- | --- | --- | --- | --- | --- |
| Temperature | 95°C | 95°C | 60°C | 72°C | 72°C | 4°C |
| Duration | 3 min | 0.25 min | 0.25 min | 0.25 min | 1 min | ∞ |
| Cycles | 1 | 35 |  |  | 1 |  |

### Trans-cardial perfusion and RNAscope fluorescence in situ hybridzation (FISH)

For RNAscope FISH, mice aged P30 were anesthetized with a sodium pentobarbital solution (Euthanasia-III Solution, Med-Pharmex, Pomona, CA; 390 mg/ml, 3.9 g/Kg). The mice were perfused through the ascending aorta with 20 ml PBS and 20 ml 4% PFA/PBS at 4°C at 8 ml/min. The brains were removed, post-fixed with 4% PFA/PBS overnight at 4°C, cryo-protected in 30% sucrose PBS at 4°C for 48 hours and stored in PBS for 24 hours. To prepare coronal sections for RNAscope, we separated the two hemispheres, embedded them in 4% w/v agar/PBS and prepared 40 µm thick slices using a vibrating blade microtome (VT1200S, Leica Microsystems, Buffalo Grove, IL). The slices were post-fixed in 4% PFA/PBS for 30 min at room temperature (RT) and mounted onto Superfrost plus microscope slides. Mounted slices were used for fluorescence in situ hybridization (FISH) using an RNAscope multiplex fluorescent assay (Advanced Cell Diagnostics, Newark, CA) according to manufacturer instructions, using Mm-Drd1-C1, Mm-Sc1a1-C2 and Mm-Drd2-C3 RNA probes and Opal 520, 570 and 690 dyes (Akoya Biosciences, Menlo Park, CA). DAPI Fluoromount G was used as the mounting medium (Cat# 0100-02; SouthernBiotech, Birmingham, AL). The presence of mRNA transcripts was assessed using a confocal microscope (Zeiss LSM710) equipped with a Plan-Apochromat 63X/1.4NA oil objective. Image size was set to 1,024 · 1,024 pixels and represented the average of 8 consecutive frames. The image analysis was performed using CellProfiler version 4.0.7 and a modified version of the Speckle Counting Pipeline (Carpenter et al. 2006). In this pipeline, we set a feature size of 20 for the EnhanceOrSuppressFeatures tool. We used Advanced Settings for the IdentifyPrimaryObjects tool. Here, the typical diameter of objects was set to 3-20 pixels, the selected thresholding method was Robust Background, and the Upper Outlier Fraction was set to 0. The size of the smoothing filter for declumping was set to 4. The minimal allowed distance to suppress local maxima was set to 4. We did not use lower-resolution images to find local maxima. The EnhanceOrSuppressFeatures, MaskImage, IdentifyPrimaryObjects and RelateObjects tools were applied for each label color (green, red, far red; i.e., 3 times).

### Stereotaxic intracranial injections and optogenetics

AAV5-EF1a-DIO-hChR2(H134R)-EYFP-WPRE-pA (University of North Carolina Vector Core, Chapel Hill, NC) was injected into D1^Cre/+^, A2A^Cre/+^, D1^Cre/+^:EAAC1^-/-^, A2A^Cre/+^:EAAC1^-/-^ mice. PHP.eB-S5E2-C1V1-eYFP (Addgene cat# 135633) was injected in D1^Cre/+^:Ai9^Tg/Tg^ mice, where it transfected ∼0.14% of all cells in the DLS. Male and female mice (P14-16) were anesthetized with isoflurane (induction: 5% in 100% O_2_ at 1-2 l/min; maintenance: 3% in 100% O_2_ at 1-2 l/min) and placed in the stereotaxic frame of a motorized drill and injection robot (Neurostar GmbH, Tübingen, Germany). After making a skin incision and thinning the skull under aseptic conditions, we injected 200 nl of the viral constructs bilaterally in the DLS using a Hamilton syringe at a rate of 100 nl/min. The injection coordinates from bregma were AP: +0.2 mm, ML: ±2.2 mm, DV: +3.0 mm. After the stereotaxic injections, the mice were returned to their home cage and used for slice physiology experiments 2-3 weeks after surgery.

### Acute slice preparation

Acute coronal slices of the mouse striatum were obtained from D1^Cre/+^:Ai9^Tg/Tg^, A2A^Cre/+^:Ai9^Tg/Tg^, D1^tdT/+^ and D1^Cre/+^:Ai9^Tg/Tg^:EAAC1^-/-^, A2A^Cre/+^:Ai9^Tg/Tg^:EAAC1^-/-^, D1^tdT/+^:EAAC1^-/-^ mice of either sex (P28-37), deeply anesthetized with isoflurane and decapitated in accordance with SUNY Albany Animal Care and Use Committee guidelines. The brain was rapidly removed and placed in ice-cold slicing solution bubbled with 95% O_2_/5% CO_2_ containing the following (in mM): 119 NaCl, 2.5 KCl, 0.5 CaCl_2_, 1.3 MgSO_4_·H_2_O, 4 MgCl_2_, 26.2 NaHCO_3_, 1 NaH_2_PO_4_, and 22 glucose, 320 mOsm, pH7.4. The slices (250 µm thick) were prepared using a vibrating blade microtome (VT1200S; Leica Microsystems, Buffalo Grove, IL). Once prepared, the slices were stored in this solution in a submersion chamber at 36°C for 30 min and at RT for at least 30 min and up to 5 hr.

### Electrophysiology, optogenetics and glutamate uncaging

Unless otherwise stated, the recording solution contained the following (in mM): 119 NaCl, 2.5 KCl, 1.2 CaCl_2_, 1 MgCl_2_, 26.2 NaHCO_3_, and 1 NaH_2_PO_4_, 22 glucose, 300 mOsm, pH7.4. In experiments performed in the absence of external Mg^2+^, the CaCl_2_ concentration was increased to 3.8 mM. We identified the DLS under bright field illumination using an upright fixed-stage microscope (BX51 WI; Olympus, Center Valley, PA). Stimulating and recording electrodes were both placed in the DLS ∼100 µm away from each other. Recordings from D1- and D2-MSNs, visually identified for their expression of the reporter protein tdTomato, were made with patch pipettes containing (in mM): 120 CsCH_3_SO_3_, 10 EGTA, 20 HEPES, 2 MgATP, 0.2 NaGTP, 5 QX-314Br, 290 mOsm, pH 7.2. Postsynaptic responses were evoked using electrical or optogenetic stimulation. Electrical stimulation was obtained by delivering constant voltage electrical pulses (50 µs) through a stimulating bipolar stainless-steel electrode (Cat# MX21AES(JD3); Frederick Haer Corporation, Bowdoin, ME). To activate ChR2-expressing fibers from MSNs or C1V1-expressing PV-INs, we used 5 ms-long light pulses generated by a SOLA-SE light engine (Lumencor, Beaverton, OR) and filtered using either a green FITC filter set (469/497/525) or a red TRITC filter set (542/570/620). The light power at the sample plane was ∼250 µW and the light pulses were delivered at intervals of 30 s using whole-field illumination through a 40X water immersion objective (LUMPLFLN40XW; Olympus, Center Valley, PA). For all electrophysiology experiments, the resistance of the recording electrode was 5-7 MOhm and was monitored throughout the experiments. Data were discarded if the resistance changed >20% during the experiment. When recording excitatory currents, picrotoxin (100 µM) was added to the recording solution to block GABA_A_ receptors. All recordings were obtained using a Multiclamp 700B amplifier and filtered at 10 KHz (Molecular Devices, San Jose, CA), converted with an 18-bit 200 kHz A/D board (HEKA, Holliston, MA), digitized at 10 KHz, and analyzed offline with custom-made software (A.S.) written in IgorPro 6.37 (Wavemetrics, Lake Oswego, OR). Tetrodotoxin (TTX; Cat# T-550) was purchased from Alomone Labs (Jerusalem, Israel). NBQX disodium salt (Cat# HB0443) and D,L-APV (Cat# HB0251), were purchased from Hello Bio (Princeton, NJ). (3S)-3-[[3-[[4-(Trifluoromethyl)benzoyl]amino]phenyl]methoxy]-L-aspartic acid, here referred to as T-TBOA (Cat# 2532), was purchased from Tocris Bio-Techne (Minneapolis, MN). D-2-Aminoadipic acid, 98% (D-AA; Cat# AAH27345MD) was purchased from Alfa Aesar (Tewksbury, MA). All other chemicals were purchased from Millipore Sigma. All recordings were performed at RT. Electrophysiological recordings were analyzed within the Igor Pro environment using custom made software (A.S.).

### Confocal imaging of biocytin filled MSNs and spine analysis

Biocytin 0.2%–0.4% (w/v) was added to the intracellular solution used to patch MSNs. Each cell was filled for at least 20 min. The slices were then fixed overnight at 4°C in 4% PFA/PBS, cryo-protected in 30% sucrose PBS, and incubated in 0.1% streptavidin-Alexa Fluor 488 or streptavidin-Alexa Fluor 647 conjugate and 0.1% Triton X-100 for 3 hours at RT. The slices were then mounted onto microscope slides using Fluoromount-G or DAPI Fluoromount-G mounting medium (Cat# 0100-01 and Cat# 0100-02, respectively; SouthernBiotech, Birmingham, AL). Confocal images were acquired using a Zeiss LSM710 inverted microscope equipped with 488 nm Ar or 633 nm HeNe laser. All images were acquired as stitched z-stacks of 4-5 frames averages (1024×1024 pixels; 1 µm z-step) using a 40X/1.4 NA or 63X/1.4NA Plan-Apochromat oil-immersion objectives. We generated 2D maximum intensity projections of biocytin-filled MSNs and traced the dendritic processes manually using Simple Neurite Traces in Fiji (https://fiji.sc/). Confocal images for spine analysis were also collected as z-stacks of 8 frame averages (1024×1024 pixels; 0.5 µm z-step; 3-5 digital zoom) using a 63X/1.4 NA Plan-Apochromat oil-immersion objective. For structural analysis, dendritic spines were classified into four groups according to their neck and head size (i.e., mushroom, thin, stubby, filopodia), using Fiji (Peters and Kaiserman-Abramof 1970; Jones and Powell 1969; Harris, Jensen, and Tsao 1992).

### NEURON model of D1-MSNs: 3D realistic morphologies of D1-MSNs

All files pertaining to the 3D realistic morphology model and simulation were uploaded to the ModelDB database (https://senselab.med.yale.edu/modeldb/, ModelDB acc.n. 267267, “Realistic Morphology” folder). 3D reconstructions of biocytin-filled D1-MSNs from EAAC1^+/+^ and EAAC1^-/-^ mice were created using the SNT plugin for Fiji (https://fiji.sc/). The dendritic arbor of these cells differed between EAAC1^+/+^ and EAAC1^-/-^ mice, consistent with data collected from 3D reconstructions of biocytin-filled D1-MSNs (**Fig. 2**). These morphologies were used to create a NEURON model that was populated with voltage-gated sodium, calcium (Ca_V_1.2, Ca_V_1.3), and potassium channels (K_AS_, K_DR_, K_IR_, K_RP_). In order to closely reproduce the main passive and active membrane properties of D1-MSNs measured experimentally, including their resting membrane potential, action potential threshold and peak (**Supp. Fig. 7-1A-D**), and repetitive firing rate in response to sustained (500 ms) somatic current injections of increasing amplitude (10-80 pA; **Supp. Fig. 7-1E-G**). Each conductance was adjusted as follows: sodium channels (2.4 S/cm^2^); voltage-gated calcium channels (Ca_V_1.2: 6.7e-6 S/cm^2^, Ca_V_1.3: 1.0e-4 S/cm^2^) and potassium channels (K_AS_: 4.0e-5 S/cm^2^, K_DR_: 5.0e-3 S/cm^2^, K_IR_: 1.0e-4 S/cm^2^, K_RP_: 2.0e-4 S/cm^2^). Voltage-gated sodium channels were placed in the axon initial segment (a 50 µm axonal segment with a diameter of 0.5 µm). All voltage-gated calcium and potassium channels were distributed throughout the plasma membrane of the soma and the dendrites. Passive leak channels were present throughout the plasma membrane.

We did not make assumptions on the location of the synaptic inputs active during our electrophysiology recordings, or on the potential existence of synaptic scaling mechanisms that might balance for passive attenuation of distal synaptic inputs, as this has not been documented for D1-MSNs (Nicholson et al. 2006; Magee and Cook 2000). Instead, we assumed that all inputs onto D1-MSNs are of equal strength, regardless of which portion of the dendrite they target. This assumption was important to set the synaptic weights of E/I inputs to ensure the model reproduced the amplitude of mEPSCs and mIPSCS recorded experimentally from D1-MSNs. To accomplish this, each simulation was repeated 100 times, each time randomizing the location of either one excitatory or inhibitory input (**Supp. Fig. 7-2A, Supp. Fig. 7-4A**). We then averaged the somatic mPSCs and compared their amplitude and kinetics with those of mPSCs recorded from D1-MSNs in EAAC1^+/+^ mice. We performed an initial adjustment of the peak conductance of the synaptic inputs in the model (AMPA: 7.2e-4 µS, GABA_A_: 3.6e-4 µS) to match the mPSC amplitude recorded from experiments.

We modeled mPSCs decay using a mono-exponential function, whereby *decay = A_l_e^-t/^*τ_1_. We set the amplitude (A_1_) and decay time constant (τ_1_) of mPSCs in the model to match that of A_1_ and τ_1_ measured across experiments (**Supp. Fig. 7-2C, D, Supp. Fig. 7-4C, D**). We tested a range of combinations of A_1_ and τ_1_ that could reproduce either the mPSC amplitude or decay time (**Supp. Fig. 7-2B; Supp. Fig. 7-4B**), to find the one that would accurately reproduce both the amplitude and kinetics of mPSCs recorded experimentally in D1-MSNs in EAAC1^+/+^ and EAAC1^-/-^ mice (**Supp. Fig. 7-2E, F; Supp. Fig. 7-4E, F**). In the case of NMDA EPSCs, we did not make a direct comparison between the model and experimental NMDA mEPSCs, as these events are particularly small in D1-MSNs, and technically challenging to record. Instead, we relied on knowledge of the AMPA mEPSC amplitude at V_hold_=-70 mV (**Fig. 3**), the reversal potential for glutamatergic currents (0 mV), and the experimentally measured value for the NMDA/AMPA ratio (EAAC1^-/-^ 0.57±0.14, n=9; EAAC1^+/+^ 0.68±0.14, n=11; p=0.57; **Supp. Fig. 3-1A, B**) to predict the NMDA mEPSC amplitude at V_hold_=40 mV. The estimated NMDA mEPSC amplitude was calculated as the product of the AMPA mEPSC amplitude at V_hold_=-70 mV, the NMDA/AMPA ratio, and the ratio of the NMDA and AMPA driving force (EAAC1^+/+^ 15.0 pA · 0.68 · 40 mV/70 mV=5.8 pA; NMDA mEPSC EAAC1^-/-^ 19.3 pA · 0.57 · 40 mV/70 mV=6.3 pA). This estimated mEPSC amplitude was used for the initial adjustment of the peak conductance of the synaptic NMDA input in the model (NMDA: 6.8e-5 µS). The estimated NMDA mEPSC decay matched that of evoked NMDA EPSCs recorded experimentally (**Supp. Fig. 7-3**). We then determined the values for A_1_ and τ_1_ for synaptic NMDA inputs in our model (**Supp. Fig. 7-3C, D**) using the same method previously mentioned for synaptic AMPA and GABA inputs. Together, these parameter-constraints allowed the compartmental model to reproduce the larger AMPA mEPSCs amplitude (**Fig. 3A- D**), similar NMDA/AMPA ratio (**Supp. Fig. 3-1**), and smaller GABA mIPSC amplitude and decay time in D1-MSNs from EAAC1^-/-^ mice (**Fig. 4**).

The models did not account for differences in tonic inhibition between D1-MSNs of EAAC1^+/+^ and EAAC1^-/-^ mice, because we verified experimentally that the tonic GABA currents in these cells are similar (EAAC1^+/+^: 4.0±1.4 pA, n=5; EAAC1^-/-^: 5.8±5.9 pA, n=7; p=0.78). These currents were measured as the picrotoxin-sensitive currents measured in cells held at −70 mV. Similar results were obtained when comparing the tonic current density, obtained by dividing the tonic currents by the capacitance of each cell (EAAC1^+/+^: 0.056±0.019 pA/pF, n=5; EAAC1^-/-^: 0.052±0.081 pA/pF, n=7; p=0.97).

### Behavioral analysis

We performed our behavioral analysis on C57BL/6NJ wild type and EAAC1^-/-^ mice of either sex aged P28-37, maintained on as 12 h:12 h L/D cycle. Three days prior to training, and for the whole duration of the training and testing sessions, the drinking water in the home cage was supplemented with 2% citric acid (w/v). Each behavioral session lasted 5 min. During each one of the ten training sessions, a 70 µl water reward was delivered in response to a lever press, and the timing of each lever press and reward was monitored using custom made MATLAB code and a B-pod state machine (Sanworks, Rochester, NY). The reward probability was set to P_rew_=0.5‖0.9. At the end of the training period, each mouse was subject to 10 training sessions with P_rew_=0.5 and 10 training sessions with P_rew_=0.9. During the one final testing session, we changed the reward probability from P_rew_=0.5 to P_rew_=0.9 at one of the following switching time intervals (5, 10, 15, 30, 50, 75 s). The number and time distribution of lever presses and rewards for each mouse was analyzed in IgorPro 6.37 (Wavemetrics, Lake Oswego, OR).

### Quantification and statistical analysis

Data averages are presented as mean ± SEM, unless indicated otherwise in the text and figure legends. Statistical significance was determined by Student’s paired or unpaired t-test, one-way ANOVAs, or two-way repeated measures ANOVAs followed by Bonferroni post hoc tests were performed as indicated in the text. Differences were considered significant at p<0.05 (*p<0.05; **p< 0.01; ***p<0.001).

## Supplementary Figure Legends

**Supplementary Figure 3-1.**
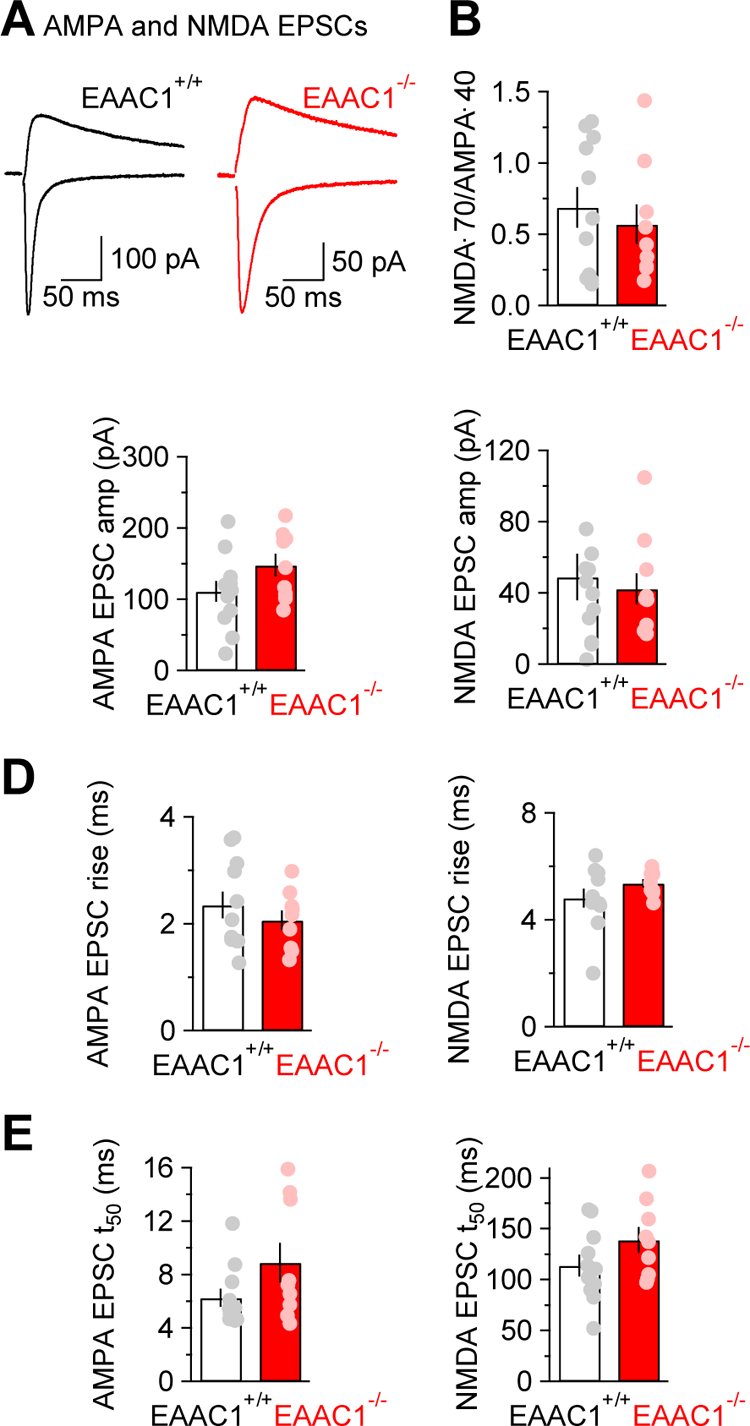
NMDA/AMPA ratio in striatal MSNs. **(A)** *Left,* Example of AMPA EPSCs recorded at V_hold_=-70 mV and NMDA EPSCs recorded at V_hold_=40 mV in a D1-MSN from an EAAC1^+/+^ mouse. Each trace is the average of 20 consecutive trials. *Right,* as in the left panel, for an EAAC1^-/-^ mouse. **(B-E)** Summary of the NMDA/AMPA ratio weighted by the driving force of the currents mediated by these receptors (B), evoked AMPA and NMDA EPSC amplitude (C), 20-80% rise time (D) and 50% decay time (E) measured in D1-MSNs of EAAC1^+/+^ (n=11) and EAAC1^-/-^ mice (n=8).

**Supplementary Figure 6-1.**
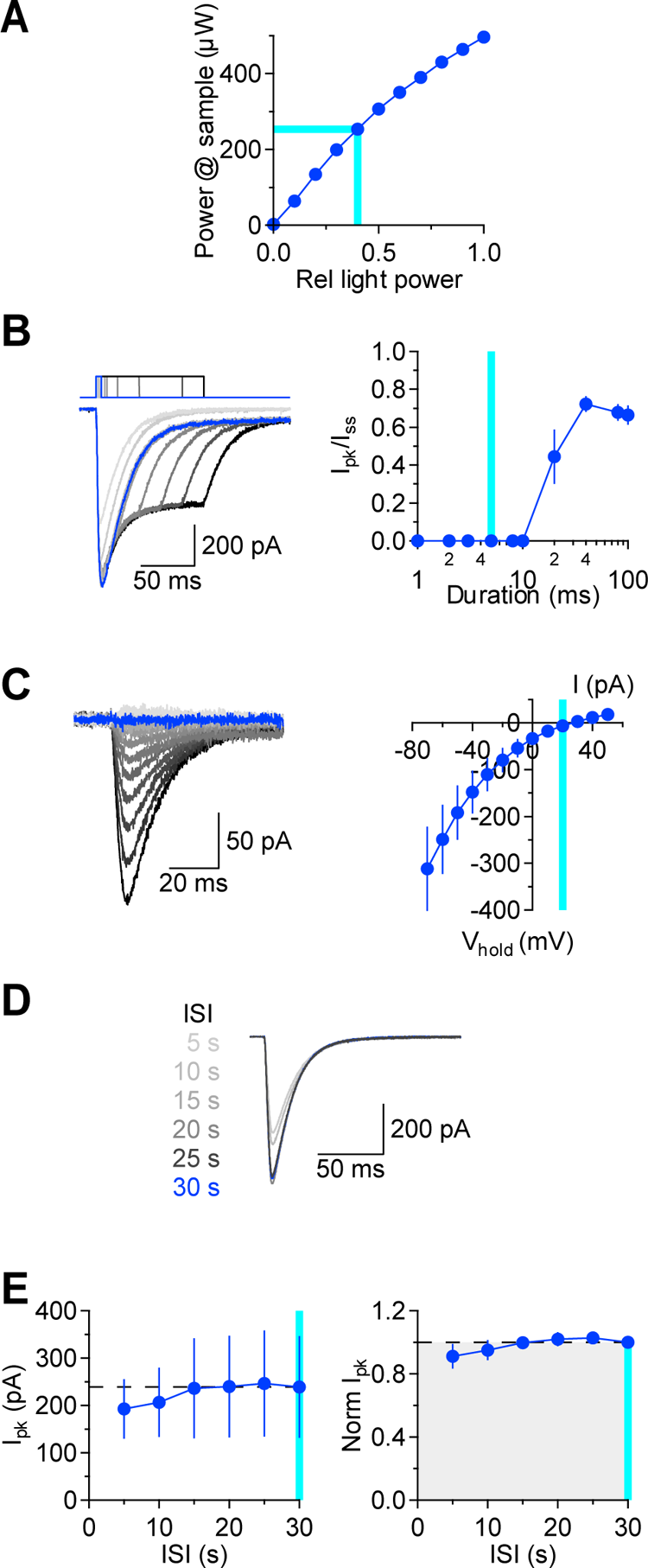
Optimization of stimulation parameters for optogenetic stimulation. **(A)** Power output of the light source used for the slice optogenetics experiments. Our experiments were performed using stimulation intensities leading ∼250 µW power at the sample plane, measured using a power meter tuned to 488 nm. **(B)** *Left,* ChR2-mediated photocurrents recorded from MSNs in EAAC1^+/+^ mice at V_hold_=-70 mV, in response to light pulses of increasing duration (1-100 ms). Each trace represents the average of 20 consecutive trials, acquired at 30 s inter-pulse intervals. The current evoked by a 5 ms-long stimulation, which we used in the rest of our experiments, is highlighted in blue. *Right,* Quantification of the amplitude ratio between the peak and steady state values of the ChR2-mediated photocurrents. Light pulses of 5 ms duration (*cyan band*) did not evoke a steady-state current (n=9). **(C)** *Left,* Example of ChR2-mediated photocurrents recorded at different holding potentials. *Right,* I/V profile of ChR2-mediated photocurrents, with an estimated reversal potential of 20 mV (*cyan band*; n=7). **(D)** *Left,* Example of ChR2 photocurrents recorded using different inter-stimulus intervals (ISIs). **(E)** *Left,* Peak amplitude of ChR2-mediated photocurrents recorded at different ISIs. No rundown was detected using 30 s ISIs, which is the value that we used in all other experiments presented here (*cyan band*; n=7). *Right,* As in the left panel, with results normalized by peak amplitude of ChR2 photocurrents recorded with ISI=30 s.

**Supplementary Figure 7-1.**
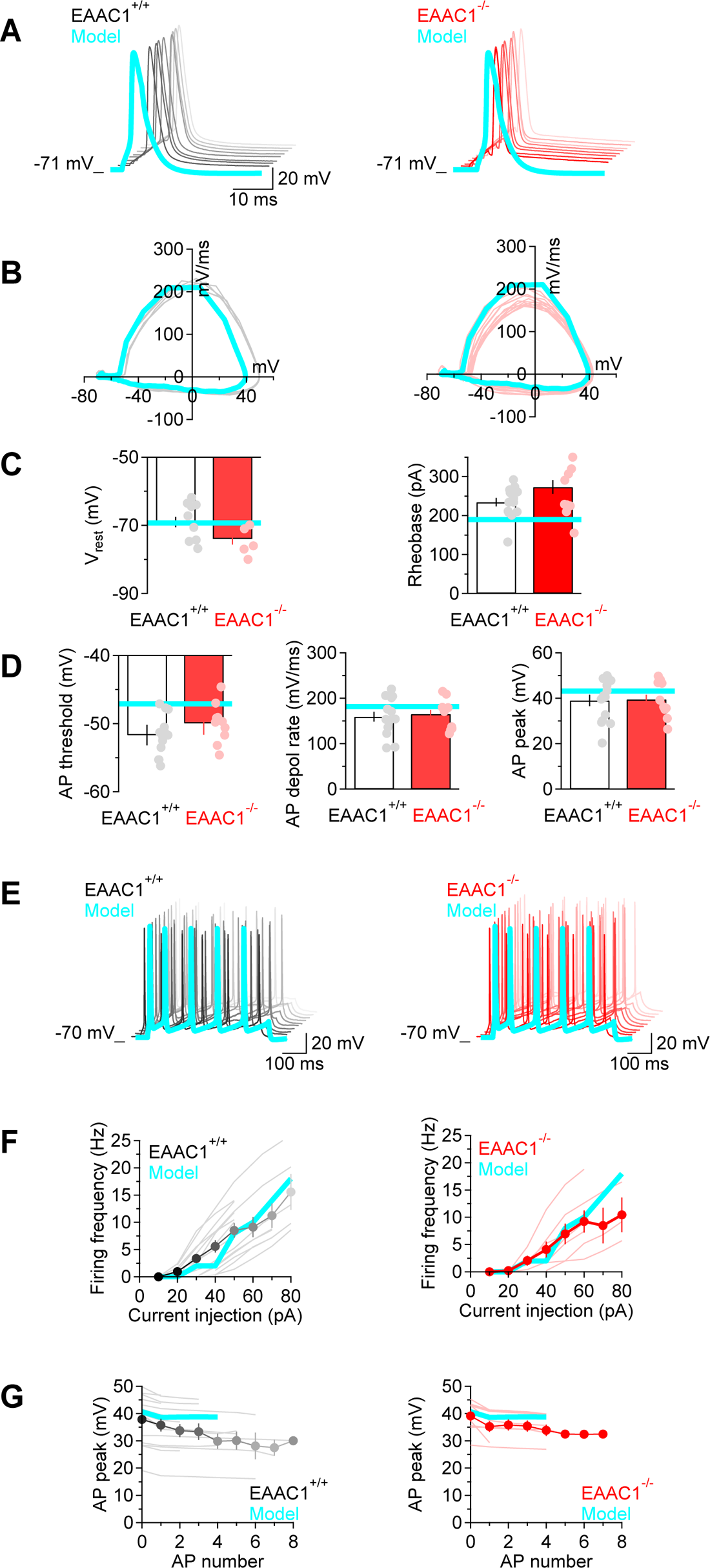
Optimization of ball-and-stick NEURON model of D1-MSN excitability. **(A)** *Left,* Example of action potentials evoked by brief somatic depolarizing steps delivered to a D1-MSN from EAAC1^+/+^ mice (150 pA · 5 ms). The action potential evoked by the NEURON model is superimposed to the traces (*cyan*). *Right,* As in the left panel, for a D1-MSN of EAAC1^-/-^ mice (150 pA · 5 ms). **(B)** Phase plots of the action potentials recorded in the experiments shown in A (*gray* and *pink* for EAAC1^+/+^ and EAAC1^-/-^ mice, respectively) and in the NEURON model (*cyan*). **(C)** *Left,* Values of the resting membrane potential measured experimentally in D1-MSNs of EAAC1^+/+^ (n=12) and EAAC1^-/-^ mice (n=9), and used in the NEURON model (*cyan line*). *Right,* As in the left panel, for the rheobase levels in D1-MSNs of EAAC1^+/+^ (n=14) and EAAC1^-/-^ (n=9). **(D)** As in C, for the action potential threshold (*left*), peak (*center*) and depolarization rate (*right*). **(E)** *Left,* Example of trains of action potentials evoked by prolonged somatic depolarizing steps delivered to a D1-MSN from EAAC1^+/+^ mice (50 pA · 500 ms). The train of action potentials evoked by the NEURON model is superimposed to the traces (*cyan*). *Right,* As in the left panel, for a D1-MSN from EAAC1^-/-^ mice (50 pA · 500 ms). **(F)** *Left,* Frequency/current (f/I) plots obtained using 500 ms somatic current step injections in D1-MSNs from EAAC1^+/+^ mice (n=14). *Right,* As in the left panel, for D1-MSNs from EAAC1^-/-^ mice (n=9). **(G)** Average peak of each action potential in a train evoked using 50 pA · 500 ms somatic current injections. Data represent mean ± SEM.

**Supplementary Figure 7-2.** Optimization of synaptic weight for excitatory synapses containing AMPA receptors, using a NEURON model of D1-MSNs with realistic morphology. **(A)** Schematic representation of the ball-and-stick NEURON compartmental model of D1-MSNs, with one AMPA input (*green*) located at varying distances from the soma. The location of the input was selected randomly and results of 100 simulations were averaged to obtain the data shown in panels B-D. **(B)** The contour plots represent an overlay of those described in panels C,D. The dots represent the only combination of A_1_ and τ_1_ values (where *decay = A_l_e^-t/n^*), which allow our model to capture the experimentally detected differences in AMPA mEPSC amplitude and decay time. **(C)** The colored matrix represents the peak normalized AMPA mEPSC amplitude obtained using different combinations of A_1_ and τ_1_. The contour plots describe the infinite number of possible solutions for currents recorded from our ball-and-stick model of D1-MSNs in EAAC1^+/+^ (*black curve*) and EAAC1^-/-^ mice (*red dashed curve*). The red dashed contour plot is obtained setting the normalized mEPSC amplitude to 1.35, as indicated above the plot. **(D)** The colored matrix represents the normalized AMPA mEPSC 50% decay time obtained using different combinations of A_1_ and τ_1_. The contour plots describe the infinite number of possible solutions for currents recorded from our ball-and-stick model of D1-MSNs in EAAC1^+/+^ (*black line*) and EAAC1^-/-^ mice (*red dashed line*). The red dashed contour plot is obtained setting the normalized mEPSC t_50_ to 1.01, as indicated above the plot. **(E)** *Left,* overlay of AMPA mEPSCs recorded from D1-MSNs in EAAC1^+/+^ (*black*) and EAAC1^-/-^ mice (*red*). Each trace represents the average of mEPSCs recorded from EAAC1^+/+^ (n=12) and EAAC1^-/-^ D1-MSNs (n=10). *Right,* overlay of modeled AMPA mEPSCs in D1-MSNs from EAAC1^+/+^ (*black*) and EAAC1^-/-^ mice (*red*). **(F)** As in E, after peak normalization.

**Supplementary Figure 7-3.**
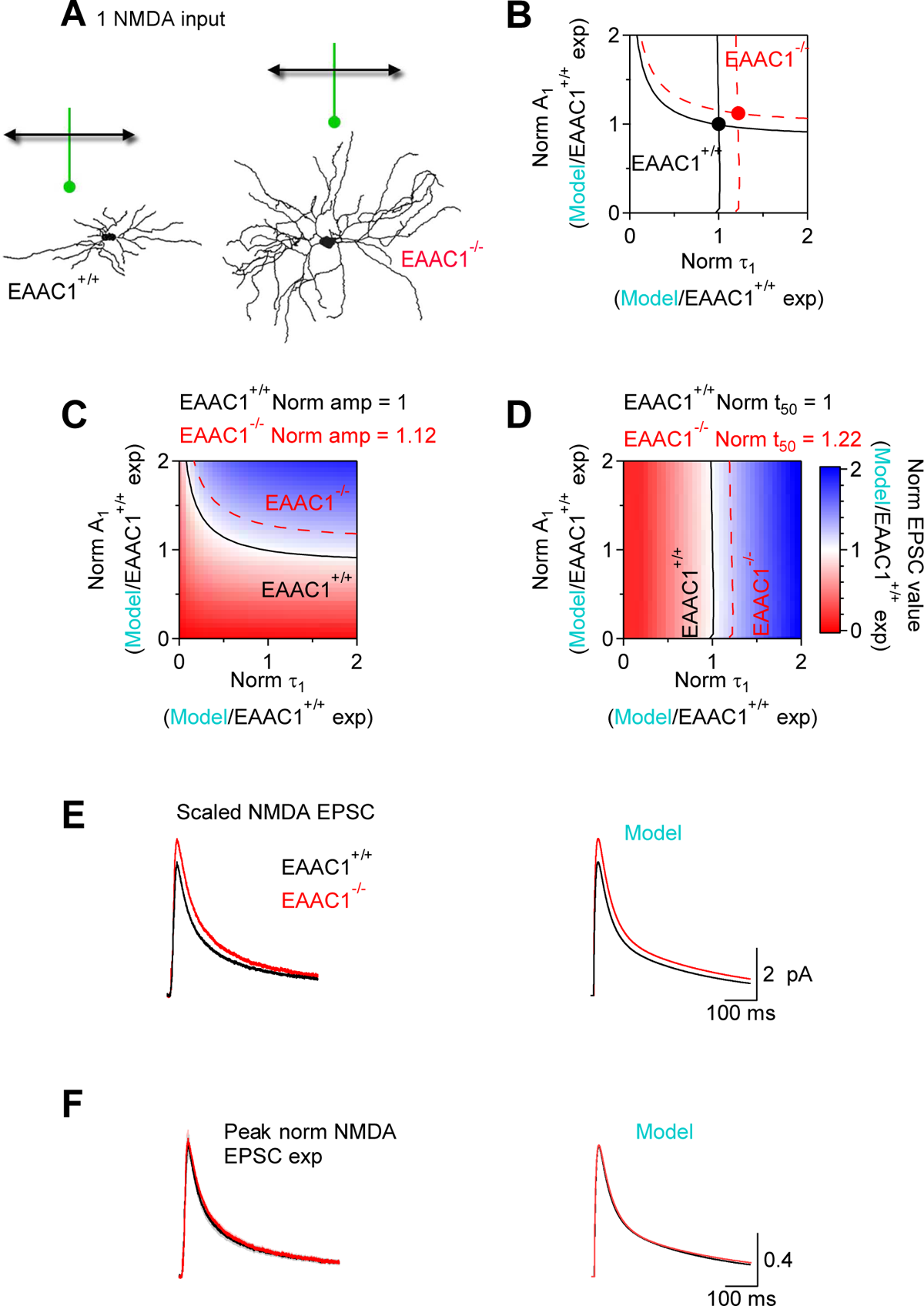
Optimization of synaptic weight for excitatory synapses containing NMDA receptors, using a NEURON model of D1-MSNs with realistic morphology. **(A)** Schematic representation of the ball-and-stick NEURON compartmental model of D1-MSNs, with one NMDA input (*green*) located at varying distances from the soma. The location of the input was selected randomly and results of 100 simulations were averaged to obtain the results shown in panels B-D. **(B)** The contour plots represent an overlay of those described in panels C, D. The dots represent the only combination of A_1_ and τ_1_ values that allow our model to capture the experimentally detected differences in NMDA EPSC amplitude and decay time, after their amplitude was scaled based on the amplitude of AMPA mEPSCs and the NMDA/AMPA ratio. **(C)** The colored matrix represents the peak normalized NMDA EPSC amplitude obtained using different combinations of A_1_ and τ_1_. The contour plots describe the infinite number of possible solutions for currents recorded from our ball-and-stick model of D1-MSNs in EAAC1^+/+^ (*black curve*) and EAAC1^-/-^ mice (*red dashed curve*). The red dashed contour plot is obtained setting the normalized EPSC amplitude to 1.12, as indicated above the plot. **(D)** The colored matrix represents the estimated normalized NMDA mEPSC 50% decay time obtained using different combinations of A_1_ and τ_1_. The contour plots describe the infinite number of possible solutions for currents recorded from our ball-and-stick model of D1-MSNs in EAAC1^+/+^ (*black line*) and EAAC1^-/-^ mice (*red dashed line*). The red dashed contour plot is obtained setting the normalized mEPSC t_50_ to 1.01, as indicated above the plot. **(E)** *Left,* overlay of NMDA mEPSCs estimated from recordings of evoked NMDA EPSC currents and AMPA/NMDA ratios in D1-MSNs in EAAC1^+/+^ (*black*) and EAAC1^-/-^ mice (*red*). Each trace represents the average of scaled NMDA EPSCs recorded from EAAC1^+/+^ (n=11) and EAAC1^-/-^ D1-MSNs (n=9). *Right,* overlay of modeled NMDA mEPSCs in D1-MSNs from EAAC1^+/+^ (*black*) and EAAC1^-/-^ mice (*red*). **(F)** As in E, after peak normalization.

**Supplementary Figure 7-4.**
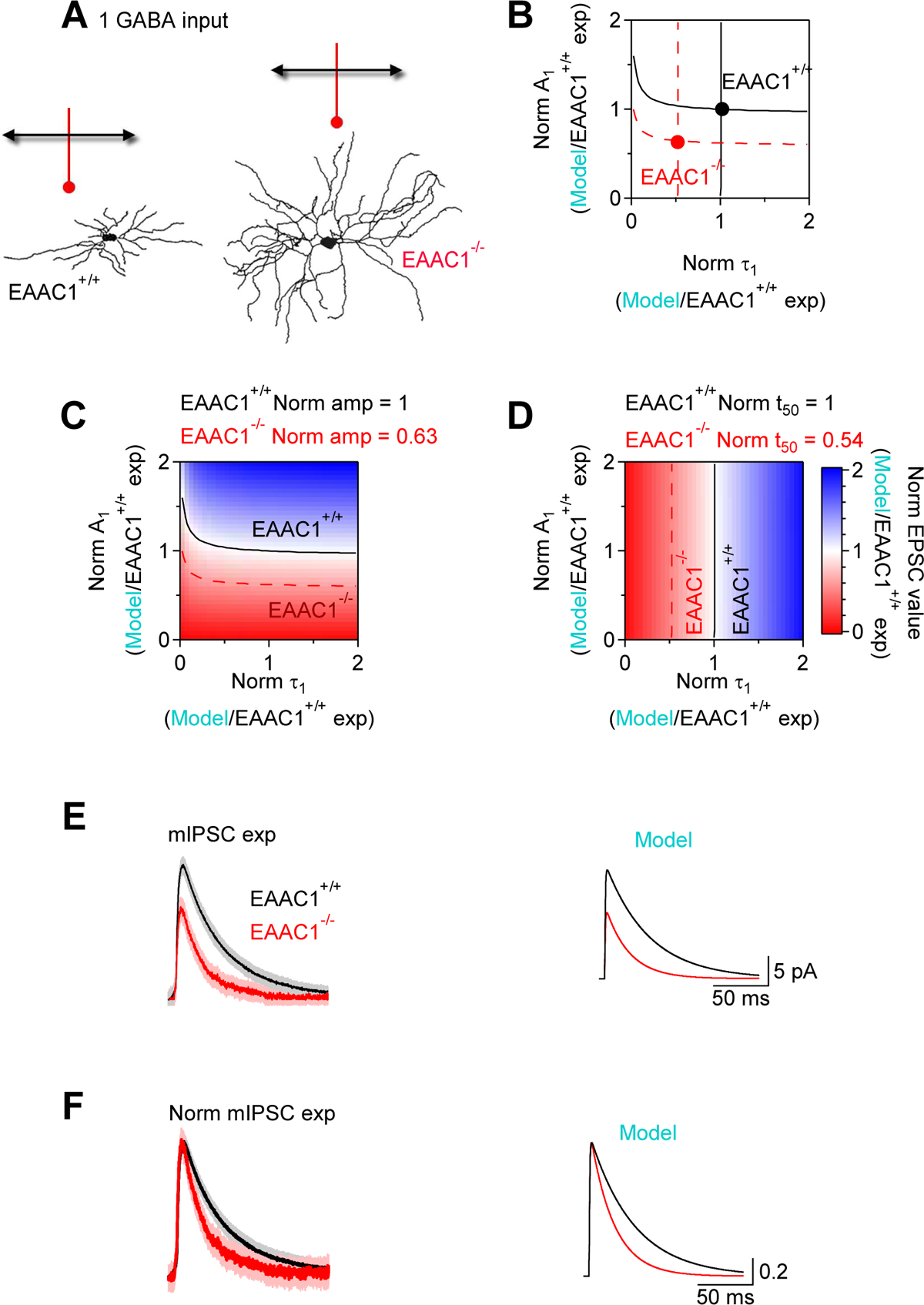
Optimization of synaptic weight for inhibitory synapses containing GABA_A_ receptors, using a NEURON model of D1-MSNs with realistic morphology. **(A)** Schematic representation of the ball-and-stick NEURON compartmental model of D1-MSNs, with one GABA input (*red*) located at varying distances from the soma. The location of the input was selected randomly and results of 100 simulations were averaged to obtain the results shown in panels B-D. **(B)** The contour plots represent an overlay of those described in panels C, D. The dots represent the only combination of A_1_ and τ_1_ values that allow our model to capture the experimentally detected differences in GABA mIPSC amplitude and decay time. **(C)** The colored matrix represents the peak normalized GABA mIPSC amplitude obtained using different combinations of A_1_ and τ_1_. The contour plots describe the infinite number of possible solutions for currents recorded from our ball-and-stick model of D1-MSNs in EAAC1^+/+^ (*black curve*) and EAAC1^-/-^ mice (*red dashed curve*). The red dashed contour plot is obtained setting the normalized EPSC amplitude to 0.63, as indicated above the plot. **(D)** The colored matrix represents the estimated normalized NMDA mEPSC 50% decay time obtained using different combinations of A_1_ and τ_1_. The contour plots describe the infinite number of possible solutions for currents recorded from our ball-and-stick model of D1-MSNs in EAAC1^+/+^ (*black line*) and EAAC1^-/-^ mice (*red dashed line*). The red dashed contour plot is obtained setting the normalized mIPSC t_50_ to 0.54, as indicated above the plot. **(E)** *Left,* overlay of GABA mIPSCs from recordings in D1-MSNs of EAAC1^+/+^ (*black*) and EAAC1^-/-^ mice (*red*). Each trace represents the average of GABA mIPSCs recorded from 11 EAAC1^+/+^ and 10 EAAC1^-/-^ D1-MSNs. *Right,* overlay of modeled GABA mIPSCs in D1-MSNs from EAAC1^+/+^ (*black*) and EAAC1^-/-^ mice (*red*). **(F)** As in E, after peak normalization.

## Acknowledgements

This work was supported by the National Science Foundation (IOS1655365). We would like to thank B. L. Sabatini for comments on the manuscript and valuable discussions.

## Data availability

All primary data used in this work and complete statistical analyses for each figure have been deposited to the Open Science Framework (https://osf.io/dw5n7/).

## References

1. Abbott, L. F., J. A. Varela, K. Sen, and S. B. Nelson. 1997. ‘Synaptic depression and cortical gain control’, Science, 275: 220–4.

2. Abramowitz, J. S., S. Taylor, and D. McKay. 2009. ‘Obsessive-compulsive disorder’, Lancet, 374: 491–9.

3. Adler, A., S. Katabi, I. Finkes, Z. Israel, Y. Prut, and H. Bergman. 2012. ‘Temporal convergence of dynamic cell assemblies in the striato-pallidal network’, J Neurosci, 32: 2473–84.

4. Arriza, J. L., W. A. Fairman, J. I. Wadiche, G. H. Murdoch, M. P. Kavanaugh, and S. G. Amara. 1994. ‘Functional comparisons of three glutamate transporter subtypes cloned from human motor cortex’, J Neurosci, 14: 5559–69.

5. Barbera, G., B. Liang, L. Zhang, C. R. Gerfen, E. Culurciello, R. Chen, Y. Li, and D. T. Lin. 2016. ‘Spatially Compact Neural Clusters in the Dorsal Striatum Encode Locomotion Relevant Information’, Neuron, 92: 202–13.

6. Bari, A., and T. W. Robbins. 2013. ‘Inhibition and impulsivity: behavioral and neural basis of response control’, Prog Neurobiol, 108: 44–79.

7. Bellini, S., K. E. Fleming, M. De, J. P. McCauley, M. A. Petroccione, L. Y. D’Brant, A. Tkachenko, S. Kwon, L. A. Jones, and A. Scimemi. 2018. ‘Neuronal Glutamate Transporters Control Dopaminergic Signaling and Compulsive Behaviors’, J Neurosci, 38: 937–61.

8. Benatti, B., B. Dell’Osso, C. Arici, E. Hollander, and A. C. Altamura. 2014. ‘Characterizing impulsivity profile in patients with obsessive-compulsive disorder’, Int J Psychiatry Clin Pract, 18: 156–60.

9. Berke, J. D., M. Okatan, J. Skurski, and H. B. Eichenbaum. 2004. ‘Oscillatory entrainment of striatal neurons in freely moving rats’, Neuron, 43: 883–96.

10. Boisseau, C. L., H. Thompson-Brenner, C. Caldwell-Harris, E. Pratt, T. Farchione, and D. H. Barlow. 2012. ‘Behavioral and cognitive impulsivity in obsessive-compulsive disorder and eating disorders’, Psychiatry Res, 200: 1062–6.

11. Burguiere, E., P. Monteiro, L. Mallet, G. Feng, and A. M. Graybiel. 2015. ‘Striatal circuits, habits, and implications for obsessive-compulsive disorder’, Curr Opin Neurobiol, 30: 59–65.

12. Burke, D. A., H. G. Rotstein, and V. A. Alvarez. 2017. ‘Striatal Local Circuitry: A New Framework for Lateral Inhibition’, Neuron, 96: 267–84.

13. Carmin, C. N., P. S. Wiegartz, U. Yunus, and K. L. Gillock. 2002. ‘Treatment of late-onset OCD following basal ganglia infarct’, Depress Anxiety, 15: 87–90.

14. Carpenter, A. E., T. R. Jones, M. R. Lamprecht, C. Clarke, I. H. Kang, O. Friman, D. A. Guertin, J. H. Chang, R. A. Lindquist, J. Moffat, P. Golland, and D. M. Sabatini. 2006. ‘CellProfiler: image analysis software for identifying and quantifying cell phenotypes’, Genome Biol, 7: R100.

15. Carrillo-Reid, L., S. Hernandez-Lopez, D. Tapia, E. Galarraga, and J. Bargas. 2011. ‘Dopaminergic modulation of the striatal microcircuit: receptor-specific configuration of cell assemblies’, J Neurosci, 31: 14972–83.

16. Carrillo-Reid, L., F. Tecuapetla, D. Tapia, A. Hernandez-Cruz, E. Galarraga, R. Drucker-Colin, and J. Bargas. 2008. ‘Encoding network states by striatal cell assemblies’, J Neurophysiol, 99: 1435–50.

17. Cheng, C., G. Glover, G. Banker, and S. G. Amara. 2002. ‘A novel sorting motif in the glutamate transporter excitatory amino acid transporter 3 directs its targeting in Madin-Darby canine kidney cells and hippocampal neurons’, J Neurosci, 22: 10643–52.

18. Chiu, D. N., and C. E. Jahr. 2017. ‘Extracellular Glutamate in the Nucleus Accumbens Is Nanomolar in Both Synaptic and Non-synaptic Compartments’, Cell Rep, 18: 2576–83.

19. Clements, J. D., R. A. Lester, G. Tong, C. E. Jahr, and G. L. Westbrook. 1992. ‘The time course of glutamate in the synaptic cleft’, Science, 258: 1498–501.

20. Coleman, P. D., and A. H. Riesen. 1968. ‘Evironmental effects on cortical dendritic fields. I. Rearing in the dark’, J Anat, 102: 363–74.

21. Conti, F., S. DeBiasi, A. Minelli, J. D. Rothstein, and M. Melone. 1998. ‘EAAC1, a high-affinity glutamate tranporter, is localized to astrocytes and gabaergic neurons besides pyramidal cells in the rat cerebral cortex’, Cereb Cortex, 8: 108–16.

22. Cui, G., S. B. Jun, X. Jin, M. D. Pham, S. S. Vogel, D. M. Lovinger, and R. M. Costa. 2013. ‘Concurrent activation of striatal direct and indirect pathways during action initiation’, Nature, 494: 238–42.

23. Czubayko, U., and D. Plenz. 2002. ‘Fast synaptic transmission between striatal spiny projection neurons’, Proc Natl Acad Sci U S A, 99: 15764–9.

24. Delgado-Acevedo, C., S. F. Estay, A. K. Radke, A. Sengupta, A. P. Escobar, F. Henriquez-Belmar, C. A. Reyes, V. Haro-Acuna, E. Utreras, R. Sotomayor-Zarate, A. Cho, J. R. Wendland, A. B. Kulkarni, A. Holmes, D. L. Murphy, A. E. Chavez, and P. R. Moya. 2019. ‘Behavioral and synaptic alterations relevant to obsessive-compulsive disorder in mice with increased EAAT3 expression’, Neuropsychopharmacology, 44: 1163–73.

25. Diamond, J. S. 2001. ‘Neuronal glutamate transporters limit activation of NMDA receptors by neurotransmitter spillover on CA1 pyramidal cells’, J Neurosci, 21: 8328–38.

26. Dobbs, L. K., A. R. Kaplan, J. C. Lemos, A. Matsui, M. Rubinstein, and V. A. Alvarez. 2016. ‘Dopamine Regulation of Lateral Inhibition between Striatal Neurons Gates the Stimulant Actions of Cocaine’, Neuron, 90: 1100–13.

27. Escobar, A. P., J. Martinez-Pinto, F. Silva-Olivares, R. Sotomayor-Zarate, and P. R. Moya. 2021. ‘Altered Grooming Syntax and Amphetamine-Induced Dopamine Release in EAAT3 Overexpressing Mice’, Front Cell Neurosci, 15: 661478.

28. Ettelt, S., S. Ruhrmann, S. Barnow, F. Buthz, A. Hochrein, K. Meyer, S. Kraft, C. Reck, R. Pukrop, J. Klosterkotter, P. Falkai, W. Maier, M. Wagner, H. J. Freyberger, and H. J. Grabe. 2007. ‘Impulsiveness in obsessive-compulsive disorder: results from a family study’, Acta Psychiatr Scand, 115: 41–7.

29. Freeze, B. S., A. V. Kravitz, N. Hammack, J. D. Berke, and A. C. Kreitzer. 2013. ‘Control of basal ganglia output by direct and indirect pathway projection neurons’, J Neurosci, 33: 18531–9.

30. Friedlander, M. J., L. R. Stanford, and S. M. Sherman. 1982. ‘Effects of monocular deprivation on the structure-function relationship of individual neurons in the cat’s lateral geniculate nucleus’, J Neurosci, 2: 321–30.

31. Gasso, P., A. E. Ortiz, S. Mas, A. Morer, A. Calvo, N. Bargallo, A. Lafuente, and L. Lazaro. 2015. ‘Association between genetic variants related to glutamatergic, dopaminergic and neurodevelopment pathways and white matter microstructure in child and adolescent patients with obsessive-compulsive disorder’, J Affect Disord, 186: 284–92.

32. Gilbert, A. R., M. S. Keshavan, V. Diwadkar, J. Nutche, F. Macmaster, P. C. Easter, C. J. Buhagiar, and D. R. Rosenberg. 2008. ‘Gray matter differences between pediatric obsessive-compulsive disorder patients and high-risk siblings: a preliminary voxel-based morphometry study’, Neurosci Lett, 435: 45–50.

33. Gillan, C. M., M. Papmeyer, S. Morein-Zamir, B. J. Sahakian, N. A. Fineberg, T. W. Robbins, and S. de Wit. 2011. ‘Disruption in the balance between goal-directed behavior and habit learning in obsessive-compulsive disorder’, Am J Psychiatry, 168: 718–26.

34. Gong, S., M. Doughty, C. R. Harbaugh, A. Cummins, M. E. Hatten, N. Heintz, and C. R. Gerfen. 2007. ‘Targeting Cre recombinase to specific neuron populations with bacterial artificial chromosome constructs’, J Neurosci, 27: 9817–23.

35. Gong, S., C. Zheng, M. L. Doughty, K. Losos, N. Didkovsky, U. B. Schambra, N. J. Nowak, A. Joyner, G. Leblanc, M. E. Hatten, and N. Heintz. 2003. ‘A gene expression atlas of the central nervous system based on bacterial artificial chromosomes’, Nature, 425: 917–25.

36. Grassi, G., S. Pallanti, L. Righi, M. Figee, M. Mantione, D. Denys, D. Piccagliani, A. Rossi, and P. Stratta. 2015. ‘Think twice: Impulsivity and decision making in obsessive-compulsive disorder’, J Behav Addict, 4: 263–72.

37. Gremel, C. M., and R. M. Costa. 2013. ‘Orbitofrontal and striatal circuits dynamically encode the shift between goal-directed and habitual actions’, Nat Commun, 4: 2264.

38. Grewer, C., N. Watzke, M. Wiessner, and T. Rauen. 2000. ‘Glutamate translocation of the neuronal glutamate transporter EAAC1 occurs within milliseconds’, Proc Natl Acad Sci U S A, 97: 9706–11.

39. Guehl, D., A. Benazzouz, B. Aouizerate, E. Cuny, J. Y. Rotge, A. Rougier, J. Tignol, B. Bioulac, and P. Burbaud. 2008. ‘Neuronal correlates of obsessions in the caudate nucleus’, Biol Psychiatry, 63: 557–62.

40. Guillery, R. W. 1973. ‘Quantitative studies of transneuronal atrophy in the dorsal lateral geniculate nucleus of cats and kittens’, J Comp Neurol, 149: 423–38.

41. Guo, Q., D. Wang, X. He, Q. Feng, R. Lin, F. Xu, L. Fu, and M. Luo. 2015. ‘Whole-brain mapping of inputs to projection neurons and cholinergic interneurons in the dorsal striatum’, PLoS One, 10: e0123381.

42. Harris, K. M., F. E. Jensen, and B. Tsao. 1992. ‘Three-dimensional structure of dendritic spines and synapses in rat hippocampus (CA1) at postnatal day 15 and adult ages: implications for the maturation of synaptic physiology and long-term potentiation’, J Neurosci, 12: 2685–705.

43. He, Y., W. G. Janssen, J. D. Rothstein, and J. H. Morrison. 2000. ‘Differential synaptic localization of the glutamate transporter EAAC1 and glutamate receptor subunit GluR2 in the rat hippocampus’, J Comp Neurol, 418: 255–69.

44. Herman, M. A., and C. E. Jahr. 2007. ‘Extracellular glutamate concentration in hippocampal slice’, J Neurosci, 27: 9736–41.

45. Higley, M. J., and B. L. Sabatini. 2010. ‘Competitive regulation of synaptic Ca2+ influx by D2 dopamine and A2A adenosine receptors’, Nat Neurosci, 13: 958–66.

46. Hines, M. L., and N. T. Carnevale. 1997. ‘The NEURON simulation environment’, Neural Comput, 9: 1179–209.

47. Holmseth, S., Y. Dehnes, Y. H. Huang, V. V. Follin-Arbelet, N. J. Grutle, M. N. Mylonakou, C. Plachez, Y. Zhou, D. N. Furness, D. E. Bergles, K. P. Lehre, and N. C. Danbolt. 2012. ‘The density of EAAC1 (EAAT3) glutamate transporters expressed by neurons in the mammalian CNS’, J Neurosci, 32: 6000–13.

48. Horvath, P. M., M. K. Piazza, L. M. Monteggia, and E. T. Kavalali. 2020. ‘Spontaneous and evoked neurotransmission are partially segregated at inhibitory synapses’, Elife, 9.

49. Huerta-Ocampo, I., J. Mena-Segovia, and J. P. Bolam. 2014. ‘Convergence of cortical and thalamic input to direct and indirect pathway medium spiny neurons in the striatum’, Brain Struct Funct, 219: 1787–800.

50. Humphries, M. D., and T. J. Prescott. 2010. ‘The ventral basal ganglia, a selection mechanism at the crossroads of space, strategy, and reward’, Prog Neurobiol, 90: 385–417.

51. Humphries, M. D., R. Wood, and K. Gurney. 2009. ‘Dopamine-modulated dynamic cell assemblies generated by the GABAergic striatal microcircuit’, Neural Netw, 22: 1174–88.

52. Humphries, M. D., R. Wood, and K. Gurney. 2010. ‘Reconstructing the three-dimensional GABAergic microcircuit of the striatum’, PLoS Comput Biol, 6: e1001011.

53. Hunnicutt, B. J., B. C. Jongbloets, W. T. Birdsong, K. J. Gertz, H. Zhong, and T. Mao. 2016. ‘A comprehensive excitatory input map of the striatum reveals novel functional organization’, Elife, 5.

54. Jones, E. G., and T. P. Powell. 1969. ‘Morphological variations in the dendritic spines of the neocortex’, J Cell Sci, 5: 509–29.

55. Kawaguchi, Y., C. J. Wilson, and P. C. Emson. 1990. ‘Projection subtypes of rat neostriatal matrix cells revealed by intracellular injection of biocytin’, J Neurosci, 10: 3421–38.

56. Koos, T., and J. M. Tepper. 1999. ‘Inhibitory control of neostriatal projection neurons by GABAergic interneurons’, Nat Neurosci, 2: 467–72.

57. Kravitz, A. V., B. S. Freeze, P. R. Parker, K. Kay, M. T. Thwin, K. Deisseroth, and A. C. Kreitzer. 2010. ‘Regulation of parkinsonian motor behaviours by optogenetic control of basal ganglia circuitry’, Nature, 466: 622–6.

58. Kravitz, A. V., L. D. Tye, and A. C. Kreitzer. 2012. ‘Distinct roles for direct and indirect pathway striatal neurons in reinforcement’, Nat Neurosci, 15: 816–8.

59. Lobo, M. K., H. E. Covington, 3rd, D. Chaudhury, A. K. Friedman, H. Sun, D. Damez-Werno, D. M. Dietz, S. Zaman, J. W. Koo, P. J. Kennedy, E. Mouzon, M. Mogri, R. L. Neve, K. Deisseroth, M. H. Han, and E. J. Nestler. 2010. ‘Cell type-specific loss of BDNF signaling mimics optogenetic control of cocaine reward’, Science, 330: 385-90.

60. Magee, J. C., and E. P. Cook. 2000. ‘Somatic EPSP amplitude is independent of synapse location in hippocampal pyramidal neurons’, Nat Neurosci, 3: 895–903.

61. Mathews, G. C., and J. S. Diamond. 2003. ‘Neuronal glutamate uptake Contributes to GABA synthesis and inhibitory synaptic strength’, J Neurosci, 23: 2040–8.

62. McAllister, A. K. 2000. ‘Cellular and molecular mechanisms of dendrite growth’, Cereb Cortex, 10: 963–73.

63. McCauley, J. P., M. A. Petroccione, L. Y. D’Brant, G. C. Todd, N. Affinnih, J. J. Wisnoski, S. Zahid, S. Shree, A. A. Sousa, R. M. De Guzman, R. Migliore, A. Brazhe, R. D. Leapman, A. Khmaladze, A. Semyanov, D. G. Zuloaga, M. Migliore, and A. Scimemi. 2020. ‘Circadian Modulation of Neurons and Astrocytes Controls Synaptic Plasticity in Hippocampal Area CA1’, Cell Rep, 33: 108255.

64. Mehaffey, W. H., B. Doiron, L. Maler, and R. W. Turner. 2005. ‘Deterministic multiplicative gain control with active dendrites’, J Neurosci, 25: 9968–77.

65. Menzies, L., S. R. Chamberlain, A. R. Laird, S. M. Thelen, B. J. Sahakian, and E. T. Bullmore. 2008. ‘Integrating evidence from neuroimaging and neuropsychological studies of obsessive-compulsive disorder: the orbitofronto-striatal model revisited’, Neurosci Biobehav Rev, 32: 525–49.

66. Menzies, L., G. B. Williams, S. R. Chamberlain, C. Ooi, N. Fineberg, J. Suckling, B. J. Sahakian, T. W. Robbins, and E. T. Bullmore. 2008. ‘White matter abnormalities in patients with obsessive-compulsive disorder and their first-degree relatives’, Am J Psychiatry, 165: 1308–15.

67. Milad, M. R., and S. L. Rauch. 2012. ‘Obsessive-compulsive disorder: beyond segregated cortico-striatal pathways’, Trends Cogn Sci, 16: 43–51.

68. Mink, J. W. 1996. ‘The basal ganglia: focused selection and inhibition of competing motor programs’, Prog Neurobiol, 50: 381–425.

69. Mitchell, S. J., and R. A. Silver. 2003. ‘Shunting inhibition modulates neuronal gain during synaptic excitation’, Neuron, 38: 433–45.

70. Moyer, J. T., B. L. Halterman, L. H. Finkel, and J. A. Wolf. 2014. ‘Lateral and feedforward inhibition suppress asynchronous activity in a large, biophysically-detailed computational model of the striatal network’, Front Comput Neurosci, 8: 152.

71. Nicholson, D. A., R. Trana, Y. Katz, W. L. Kath, N. Spruston, and Y. Geinisman. 2006. ‘Distance-dependent differences in synapse number and AMPA receptor expression in hippocampal CA1 pyramidal neurons’, Neuron, 50: 431–42.

72. Pan, W. X., T. Mao, and J. T. Dudman. 2010. ‘Inputs to the dorsal striatum of the mouse reflect the parallel circuit architecture of the forebrain’, Front Neuroanat, 4: 147.

73. Park, M. R., J. W. Lighthall, and S. T. Kitai. 1980. ‘Recurrent inhibition in the rat neostriatum’, Brain Res, 194: 359–69.

74. Pauls, D. L., A. Abramovitch, S. L. Rauch, and D. A. Geller. 2014. ‘Obsessive-compulsive disorder: an integrative genetic and neurobiological perspective’, Nat Rev Neurosci, 15: 410–24.

75. Pavlov, I., L. P. Savtchenko, D. M. Kullmann, A. Semyanov, and M. C. Walker. 2009. ‘Outwardly rectifying tonically active GABAA receptors in pyramidal cells modulate neuronal offset, not gain’, J Neurosci, 29: 15341–50.

76. Peghini, P., J. Janzen, and W. Stoffel. 1997. ‘Glutamate transporter EAAC-1-deficient mice develop dicarboxylic aminoaciduria and behavioral abnormalities but no neurodegeneration’, EMBO J, 16: 3822–32.

77. Peng, Y. R., S. He, H. Marie, S. Y. Zeng, J. Ma, Z. J. Tan, S. Y. Lee, R. C. Malenka, and X. Yu. 2009. ‘Coordinated changes in dendritic arborization and synaptic strength during neural circuit development’, Neuron, 61: 71–84.

78. Pennartz, C. M., J. D. Berke, A. M. Graybiel, R. Ito, C. S. Lansink, M. van der Meer, A. D. Redish, K. S. Smith, and P. Voorn. 2009. ‘Corticostriatal Interactions during Learning, Memory Processing, and Decision Making’, J Neurosci, 29: 12831–8.

79. Peters, A., and I. R. Kaiserman-Abramof. 1970. ‘The small pyramidal neuron of the rat cerebral cortex. The perikaryon, dendrites and spines’, Am J Anat, 127: 321–55.

80. Ponzi, A., and J. Wickens. 2010. ‘Sequentially switching cell assemblies in random inhibitory networks of spiking neurons in the striatum’, J Neurosci, 30: 5894–911.

81. Ponzi, A., and J. Wickens. 2012. ‘Input dependent cell assembly dynamics in a model of the striatal medium spiny neuron network’, Front Syst Neurosci, 6: 6.

82. Ponzi, A., and J. R. Wickens. 2013. ‘Optimal balance of the striatal medium spiny neuron network’, PLoS Comput Biol, 9: e1002954.

83. Porton, B., B. D. Greenberg, K. Askland, L. M. Serra, J. Gesmonde, G. Rudnick, S. A. Rasmussen, and H. T. Kao. 2013. ‘Isoforms of the neuronal glutamate transporter gene, SLC1A1/EAAC1, negatively modulate glutamate uptake: relevance to obsessive-compulsive disorder’, Transl Psychiatry, 3: e259.

84. Prescott, S. A., and Y. De Koninck. 2003. ‘Gain control of firing rate by shunting inhibition: roles of synaptic noise and dendritic saturation’, Proc Natl Acad Sci U S A, 100: 2076–81.

85. Preston, R. J., G. A. Bishop, and S. T. Kitai. 1980. ‘Medium spiny neuron projection from the rat striatum: an intracellular horseradish peroxidase study’, Brain Res, 183: 253–63.

86. Rǎdulescu, A., J. Herron, C. Kennedy, and A. Scimemi. 2017. ‘Global and local excitation and inhibition shape the dynamics of the cortico-striatal-thalamo-cortical pathway’, Sci Rep, 7: 7608.

87. Rakic, P. 1975. ‘Role of cell interaction in development of dendritic patterns’, Adv Neurol, 12: 117–34.

88. Rakic, P., and R. L. Sidman. 1973. ‘Organization of cerebellar cortex secondary to deficit of granule cells in weaver mutant mice’, J Comp Neurol, 152: 133–61.

89. Rall, W. 1959. ‘Branching dendritic trees and motoneuron membrane resistivity’, Exp Neurol, 1: 491–527.

90. Redmond, L., A. H. Kashani, and A. Ghosh. 2002. ‘Calcium regulation of dendritic growth via CaM kinase IV and CREB-mediated transcription’, Neuron, 34: 999–1010.

91. Reess, T. J., O. G. Rus, D. A. Gursel, B. Schmitz-Koep, G. Wagner, G. Berberich, and K. Koch. 2018. ‘Association between hippocampus volume and symptom profiles in obsessive-compulsive disorder’, Neuroimage Clin, 17: 474–80.

92. Rothstein, J. D., M. Dykes-Hoberg, C. A. Pardo, L. A. Bristol, L. Jin, R. W. Kuncl, Y. Kanai, M. A. Hediger, Y. Wang, J. P. Schielke, and D. F. Welty. 1996. ‘Knockout of glutamate transporters reveals a major role for astroglial transport in excitotoxicity and clearance of glutamate’, Neuron, 16: 675–86.

93. Rothstein, J. D., L. Martin, A. I. Levey, M. Dykes-Hoberg, L. Jin, D. Wu, N. Nash, and R. W. Kuncl. 1994. ‘Localization of neuronal and glial glutamate transporters’, Neuron, 13: 713–25.

94. Salinas, E., and P. Thier. 2000. ‘Gain modulation: a major computational principle of the central nervous system’, Neuron, 27: 15–21.

95. Sara, Y., M. Bal, M. Adachi, L. M. Monteggia, and E. T. Kavalali. 2011. ‘Use-dependent AMPA receptor block reveals segregation of spontaneous and evoked glutamatergic neurotransmission’, J Neurosci, 31: 5378–82.

96. Schikorski, T., and C. F. Stevens. 1997. ‘Quantitative ultrastructural analysis of hippocampal excitatory synapses’, J Neurosci, 17: 5858–67.

97. Schwartz, O., and E. P. Simoncelli. 2001. ‘Natural signal statistics and sensory gain control’, Nat Neurosci, 4: 819–25.

98. Scimemi, A. 2014. ‘Structure, function, and plasticity of GABA transporters’, Front Cell Neurosci, 8: 161.

99. Scimemi, A., H. Tian, and J. S. Diamond. 2009. ‘Neuronal transporters regulate glutamate clearance, NMDA receptor activation, and synaptic plasticity in the hippocampus’, J Neurosci, 29: 14581–95.

100. Shashidharan, P., G. W. Huntley, J. M. Murray, A. Buku, T. Moran, M. J. Walsh, J. H. Morrison, and A. Plaitakis. 1997. ‘Immunohistochemical localization of the neuron-specific glutamate transporter EAAC1 (EAAT3) in rat brain and spinal cord revealed by a novel monoclonal antibody’, Brain Res, 773: 139–48.

101. Shimamoto, K., B. Lebrun, Y. Yasuda-Kamatani, M. Sakaitani, Y. Shigeri, N. Yumoto, and T. Nakajima. 1998. ‘DL-threo-beta-benzyloxyaspartate, a potent blocker of excitatory amino acid transporters’, Mol Pharmacol, 53: 195–201.

102. Shmelkov, S. V., A. Hormigo, D. Jing, C. C. Proenca, K. G. Bath, T. Milde, E. Shmelkov, J. S. Kushner, M. Baljevic, I. Dincheva, A. J. Murphy, D. M. Valenzuela, N. W. Gale, G. D. Yancopoulos, I. Ninan, F. S. Lee, and S. Rafii. 2010. ‘Slitrk5 deficiency impairs corticostriatal circuitry and leads to obsessive-compulsive-like behaviors in mice’, Nat Med, 16: 598–602, 1p following 02.

103. Sin, W. C., K. Haas, E. S. Ruthazer, and H. T. Cline. 2002. ‘Dendrite growth increased by visual activity requires NMDA receptor and Rho GTPases’, Nature, 419: 475–80.

104. Somogyi, P., J. P. Bolam, and A. D. Smith. 1981. ‘Monosynaptic cortical input and local axon collaterals of identified striatonigral neurons. A light and electron microscopic study using the Golgi-peroxidase transport-degeneration procedure’, J Comp Neurol, 195: 567–84.

105. Sotelo, C. 1975. ‘Anatomical, physiological and biochemical studies of the cerebellum from mutant mice. II. Morphological study of cerebellar cortical neurons and circuits in the weaver mouse’, Brain Res, 94: 19–44.

106. Stewart, S. E., J. A. Fagerness, J. Platko, J. W. Smoller, J. M. Scharf, C. Illmann, E. Jenike, N. Chabane, M. Leboyer, R. Delorme, M. A. Jenike, and D. L. Pauls. 2007. ‘Association of the SLC1A1 glutamate transporter gene and obsessive-compulsive disorder’, Am J Med Genet B Neuropsychiatr Genet, 144B: 1027-33.

107. Stewart, S. E., C. Mayerfeld, P. D. Arnold, J. R. Crane, C. O’Dushlaine, J. A. Fagerness, D. Yu, J. M. Scharf, E. Chan, F. Kassam, P. R. Moya, J. R. Wendland, R. Delorme, M. A. Richter, J. L. Kennedy, J. Veenstra-VanderWeele, J. Samuels, B. D. Greenberg, J. T. McCracken, J. A. Knowles, A. J. Fyer, S. L. Rauch, M. A. Riddle, M. A. Grados, O. J. Bienvenu, B. Cullen, Y. Wang, Y. Y. Shugart, J. Piacentini, S. Rasmussen, G. Nestadt, D. L. Murphy, M. A. Jenike, E. H. Cook, D. L. Pauls, G. L. Hanna, and C. A. Mathews. 2013. ‘Meta-analysis of association between obsessive-compulsive disorder and the 3’ region of neuronal glutamate transporter gene SLC1A1’, Am J Med Genet B Neuropsychiatr Genet, 162B: 367-79.

108. Stewart, S. E., D. Yu, J. M. Scharf, B. M. Neale, J. A. Fagerness, C. A. Mathews, P. D. Arnold, P. D. Evans, E. R. Gamazon, L. K. Davis, L. Osiecki, L. McGrath, S. Haddad, J. Crane, D. Hezel, C. Illman, C. Mayerfeld, A. Konkashbaev, C. Liu, A. Pluzhnikov, A. Tikhomirov, C. K. Edlund, S. L. Rauch, R. Moessner, P. Falkai, W. Maier, S. Ruhrmann, H. J. Grabe, L. Lennertz, M. Wagner, L. Bellodi, M. C. Cavallini, M. A. Richter, E. H. Cook, Jr., J. L. Kennedy, D. Rosenberg, D. J. Stein, S. M. Hemmings, C. Lochner, A. Azzam, D. A. Chavira, E. Fournier, H. Garrido, B. Sheppard, P. Umana, D. L. Murphy, J. R. Wendland, J. Veenstra-VanderWeele, D. Denys, R. Blom, D. Deforce, F. Van Nieuwerburgh, H. G. Westenberg, S. Walitza, K. Egberts, T. Renner, E. C. Miguel, C. Cappi, A. G. Hounie, M. Conceicao do Rosario, A. S. Sampaio, H. Vallada, H. Nicolini, N. Lanzagorta, B. Camarena, R. Delorme, M. Leboyer, C. N. Pato, M. T. Pato, E. Voyiaziakis, P. Heutink, D. C. Cath, D. Posthuma, J. H. Smit, J. Samuels, O. J. Bienvenu, B. Cullen, A. J. Fyer, M. A. Grados, B. D. Greenberg, J. T. McCracken, M. A. Riddle, Y. Wang, V. Coric, J. F. Leckman, M. Bloch, C. Pittenger, V. Eapen, D. W. Black, R. A. Ophoff, E. Strengman, D. Cusi, M. Turiel, F. Frau, F. Macciardi, J. R. Gibbs, M. R. Cookson, A. Singleton, J. Hardy, A. T. Crenshaw, M. A. Parkin, D. B. Mirel, D. V. Conti, S. Purcell, G. Nestadt, G. L. Hanna, M. A. Jenike, J. A. Knowles, N. Cox, and D. L. Pauls. 2013. ‘Genome-wide association study of obsessive-compulsive disorder’, Mol Psychiatry, 18: 788–98.

109. Sung, K. W., S. Choi, and D. M. Lovinger. 2001. ‘Activation of group I mGluRs is necessary for induction of long-term depression at striatal synapses’, Journal of Neurophysiology, 86: 2405–12.

110. Svoboda, K., D. W. Tank, and W. Denk. 1996. ‘Direct measurement of coupling between dendritic spines and shafts’, Science, 272: 716–9.

111. Taverna, S., E. Ilijic, and D. J. Surmeier. 2008. ‘Recurrent collateral connections of striatal medium spiny neurons are disrupted in models of Parkinson’s disease’, Journal of Neuroscience, 28: 5504–12.

112. Tecuapetla, F., T. Koos, J. M. Tepper, N. Kabbani, and M. F. Yeckel. 2009. ‘Differential dopaminergic modulation of neostriatal synaptic connections of striatopallidal axon collaterals’, J Neurosci, 29: 8977–90.

113. Tepper, J. M., T. Koos, and C. J. Wilson. 2004. ‘GABAergic microcircuits in the neostriatum’, Trends Neurosci, 27: 662–9.

114. Tepper, J. M., F. Tecuapetla, T. Koos, and O. Ibanez-Sandoval. 2010. ‘Heterogeneity and diversity of striatal GABAergic interneurons’, Front Neuroanat, 4: 150.

115. Tepper, J. M., C. J. Wilson, and T. Koos. 2008. ‘Feedforward and feedback inhibition in neostriatal GABAergic spiny neurons’, Brain Res Rev, 58: 272–81.

116. Tonnesen, J., G. Katona, B. Rozsa, and U. V. Nagerl. 2014. ‘Spine neck plasticity regulates compartmentalization of synapses’, Nat Neurosci, 17: 678–85.

117. Tsukada, S., M. Iino, Y. Takayasu, K. Shimamoto, and S. Ozawa. 2005. ‘Effects of a novel glutamate transporter blocker, (2S, 3S)-3-[3-[4-(trifluoromethyl)benzoylamino]benzyloxy]aspartate (TFB-TBOA), on activities of hippocampal neurons’, Neuropharmacology, 48: 479–91.

118. Tunstall, M. J., D. E. Oorschot, A. Kean, and J. R. Wickens. 2002. ‘Inhibitory interactions between spiny projection neurons in the rat striatum’, J Neurophysiol, 88: 1263–9.

119. Ullrich, M., M. Weber, A. M. Post, S. Popp, J. Grein, M. Zechner, H. Guerrero Gonzalez, A. Kreis, A. G. Schmitt, N. Uceyler, K. P. Lesch, and K. Schuh. 2018. ‘OCD-like behavior is caused by dysfunction of thalamo-amygdala circuits and upregulated TrkB/ERK-MAPK signaling as a result of SPRED2 deficiency’, Mol Psychiatry, 23: 444–58.

120. Veenstra-VanderWeele, J., C. L. Muller, H. Iwamoto, J. E. Sauer, W. A. Owens, C. R. Shah, J. Cohen, P. Mannangatti, T. Jessen, B. J. Thompson, R. Ye, T. M. Kerr, A. M. Carneiro, J. N. Crawley, E. Sanders-Bush, D. G. McMahon, S. Ramamoorthy, L. C. Daws, J. S. Sutcliffe, and R. D. Blakely. 2012. ‘Autism gene variant causes hyperserotonemia, serotonin receptor hypersensitivity, social impairment and repetitive behavior’, Proc Natl Acad Sci U S A, 109: 5469–74.

121. Vormstein-Schneider, D., J. D. Lin, K. A. Pelkey, R. Chittajallu, B. Guo, M. A. Arias-Garcia, K. Allaway, S. Sakopoulos, G. Schneider, O. Stevenson, J. Vergara, J. Sharma, Q. Zhang, T. P. Franken, J. Smith, L. A. Ibrahim, M. Astro KJ, E. Sabri, S. Huang, E. Favuzzi, T. Burbridge, Q. Xu, L. Guo, I. Vogel, V. Sanchez, G. A. Saldi, B. L. Gorissen, X. Yuan, K. A. Zaghloul, O. Devinsky, B. L. Sabatini, R. Batista-Brito, J. Reynolds, G. Feng, Z. Fu, C. J. McBain, G. Fishell, and J. Dimidschstein. 2020. ‘Viral manipulation of functionally distinct interneurons in mice, non-human primates and humans’, Nat Neurosci, 23: 1629–36.

122. Wadiche, J. I., S. G. Amara, and M. P. Kavanaugh. 1995. ‘Ion fluxes associated with excitatory amino acid transport’, Neuron, 15: 721–8.

123. Wadiche, J. I., J. L. Arriza, S. G. Amara, and M. P. Kavanaugh. 1995. ‘Kinetics of a human glutamate transporter’, Neuron, 14: 1019–27.

124. Wall, N. R., M. De La Parra, E. M. Callaway, and A. C. Kreitzer. 2013. ‘Differential innervation of direct- and indirect-pathway striatal projection neurons’, Neuron, 79: 347–60.

125. Wang, C. S., N. L. Chanaday, L. M. Monteggia, and E. T. Kavalali. 2022. ‘Probing the segregation of evoked and spontaneous neurotransmission via photobleaching and recovery of a fluorescent glutamate sensor’, Elife, 11.

126. Welch, J. M., J. Lu, R. M. Rodriguiz, N. C. Trotta, J. Peca, J. D. Ding, C. Feliciano, M. Chen, J. P. Adams, J. Luo, S. M. Dudek, R. J. Weinberg, N. Calakos, W. C. Wetsel, and G. Feng. 2007. ‘Cortico-striatal synaptic defects and OCD-like behaviours in Sapap3-mutant mice’, Nature, 448: 894–900.

127. Wendland, J. R., P. R. Moya, K. R. Timpano, A. P. Anavitarte, M. R. Kruse, M. G. Wheaton, R. F. Ren-Patterson, and D. L. Murphy. 2009. ‘A haplotype containing quantitative trait loci for SLC1A1 gene expression and its association with obsessive-compulsive disorder’, Arch Gen Psychiatry, 66: 408–16.

128. Wiesel, T. N., and D. H. Hubel. 1963. ‘Effects of Visual Deprivation on Morphology and Physiology of Cells in the Cats Lateral Geniculate Body’, J Neurophysiol, 26: 978–93.

129. Wilson, C. J. 2007. ‘GABAergic inhibition in the neostriatum’, Prog Brain Res, 160: 91–110.

130. Wilson, C. J., and P. M. Groves. 1980. ‘Fine structure and synaptic connections of the common spiny neuron of the rat neostriatum: a study employing intracellular inject of horseradish peroxidase’, J Comp Neurol, 194: 599–615.

131. Wood, J., and S. E. Ahmari. 2015. ‘A Framework for Understanding the Emerging Role of Corticolimbic-Ventral Striatal Networks in OCD-Associated Repetitive Behaviors’, Front Syst Neurosci, 9: 171.

132. Yim, M. Y., A. Aertsen, and A. Kumar. 2011. ‘Significance of input correlations in striatal function’, PLoS Comput Biol, 7: e1002254.

133. Yin, H. H., B. J. Knowlton, and B. W. Balleine. 2004. ‘Lesions of dorsolateral striatum preserve outcome expectancy but disrupt habit formation in instrumental learning’, Eur J Neurosci, 19: 181–9.

134. Yin, H. H., B. J. Knowlton, and B. W. Balleine. 2006. ‘Inactivation of dorsolateral striatum enhances sensitivity to changes in the action-outcome contingency in instrumental conditioning’, Behav Brain Res, 166: 189–96.

135. Yu, X., and R. C. Malenka. 2003. ‘Beta-catenin is critical for dendritic morphogenesis’, Nat Neurosci, 6: 1169–77.

136. Yung, K. K., A. D. Smith, A. I. Levey, and J. P. Bolam. 1996. ‘Synaptic connections between spiny neurons of the direct and indirect pathways in the neostriatum of the rat: evidence from dopamine receptor and neuropeptide immunostaining’, Eur J Neurosci, 8: 861–9.

137. Zike, I. D., M. O. Chohan, J. M. Kopelman, E. N. Krasnow, D. Flicker, K. M. Nautiyal, M. Bubser, C. Kellendonk, C. K. Jones, G. Stanwood, K. F. Tanaka, H. Moore, S. E. Ahmari, and J. Veenstra-VanderWeele. 2017. ‘OCD candidate gene SLC1A1/EAAT3 impacts basal ganglia-mediated activity and stereotypic behavior’, Proc Natl Acad Sci U S A, 114: 5719–24.

